# Towards Universal Cell Embeddings: Integrating Single-cell RNA-seq Datasets across Species with SATURN

**DOI:** 10.1101/2023.02.03.526939

**Authors:** Yanay Rosen, Maria Brbić, Yusuf Roohani, Kyle Swanson, Ziang Li, Jure Leskovec

**Affiliations:** Department of Computer Science, Stanford University, Stanford, CA, USA; School of Computer and Communication Sciences, Swiss Federal Institute of Technology (EPFL), Lausanne, Switzerland; Department of Biomedical Data Science, Stanford University, Stanford, CA, USA; Department of Computer Science and Technology, Tsinghua University, Beijing, China

## Abstract

Analysis of single-cell datasets generated from diverse organisms offers unprecedented opportunities to unravel fundamental evolutionary processes of conservation and diversification of cell types. However, inter-species genomic differences limit the joint analysis of cross-species datasets to homologous genes. Here, we present SATURN, a deep learning method for learning universal cell embeddings that encodes genes’ biological properties using protein language models. By coupling protein embeddings from language models with RNA expression, SATURN integrates datasets profiled from different species regardless of their genomic similarity. SATURN has a unique ability to detect functionally related genes co-expressed across species, redefining differential expression for cross-species analysis. We apply SATURN to three species whole-organism atlases and frog and zebrafish embryogenesis datasets. We show that cell embeddings learnt in SATURN can be effectively used to transfer annotations across species and identify both homologous and species-specific cell types, even across evolutionarily remote species. Finally, we use SATURN to reannotate the five species Cell Atlas of Human Trabecular Meshwork and Aqueous Outflow Structures and find evidence of potentially divergent functions between glaucoma associated genes in humans and other species.

## Introduction

Cell mapping consortia efforts have generated large-scale single-cell datasets comprising hundreds of thousands of cells with the goal of uncovering underlying cellular processes. In-depth analysis of diverse datasets generated across different species through global efforts such as the Human Cell Atlas [1, 2], the Mouse Cell Atlas [3] and the Fly Cell Atlas [4, 5] has broadened our understanding of cell biology characterizing many cell types for the first time. However, current analyses remain limited in their ability to jointly analyze datasets generated across different species. Such joint analysis offers great potential for understanding fundamental evolutionary processes such as identifying cell types that are conserved across species and identifying the corresponding gene programs that drive similarities and differences of such cell types.

A variety of linear [6] [7] and, more recently, deep learning approaches [8–10] have been developed to learn low-dimensional representations of single-cell RNA scRNA expression data (cell embeddings). However, existing methods represent genes only as columns of an RNA expression matrix and thus, do not account for the biological properties of genes. This severely limits their usability when analyzing datasets generated from different species in which only a subset of genes can be matched as one-to-one homologs. While sequence alignment methods have been explored to incorporate weighted relationships between genes across species [11], they are dependent on arbitrary alignment thresholds and do not capture remote homology. Recent advances in protein language models trained on hundreds of millions protein sequences [12–14] suggest strong potential in addressing these issues by learning informative representations of the proteins a gene encodes. This is evidenced through the remarkable ability of protein representations to encode protein structure, function, molecular properties [12] and homology [15]. However, so far, the representational power of these models has not been exploited to learn cell representations that capture functional similarity of genes.

We present SATURN (Species Alignment Through Unification of Rna and proteiNs), a deep learning approach that integrates cross-species single-cell RNA sequencing (scRNA-seq) datasets by coupling gene expression with protein embeddings generated by large protein language models. SATURN introduces a concept of macrogenes defined as groups of genes that share similar protein embeddings. The strength of associations of genes to macrogenes is learned to reflect this similarity, thereby allowing functionally-related genes as captured by the protein embeddings, to group together. SATURN is uniquely able to perform multi-species differential expression analysis revealing functionally related groups of genes co-expressed across species. By mapping single-cell datasets generated with different genes to a joint embedding space, SATURN takes major steps towards universal cell embeddings.

We apply these embeddings to diverse tasks such as integration of cross-species cell atlas datasets, discovery of species specific cell types, reannotation and cross-species label transfer, as well as identification of protein differences across species. In particular, we apply SATURN to integrate Tabula Sapiens [2], Tabula Microcebus [16], and Tabula Muris [3] cell atlas datasets, creating a mammalian cell atlas of 335, 000 cells across 9 common tissues. We further apply SATURN to integrate frog and zebrafish embryogenesis datasets [17]. Our results show that SATURN successfully transfers annotations even across evolutionarily remote species and finds homologous and species-specific cell types, outperforming existing cross-species integration methods by a large margin. Finally, we apply SATURN to reannotate the five species Cell Atlas of Human Trabecular Meshwork and Aqueous Outflow Structures [18]. We find that SATURN identifies glaucoma associated macrogenes that have potentially divergent functions across species.

## Results

### Overview of SATURN

The major challenge of cross-species integration is that different datasets have different genes that may not have common one-to-one homologs. Subsetting each species’ set of genes to the common set of one-to-one homologs leads to losing a large portion of biologically relevant genes. Increasing the number of species exacerbates this problem, since a gene must have a homolog in each species in order to be considered for integration. SATURN overcomes this problem by using large protein language models to learn cell embeddings that encode the biological meaning of genes. SATURN maps cross-species datasets in the space of functionally related genes determined by protein embeddings. SATURN’s use of protein language models allows it to represent functional similarities even between remotely homologous genes that are missed by integration methods that rely on sequence based similarity [11].

In particular, SATURN integrates scRNA-seq datasets generated from different species with different genes by mapping them to a joint low-dimensional embedding space using gene expression and protein representations. SATURN takes as input: *(i)* scRNA-seq count data from one or multiple species, *(ii)* protein embeddings generated by a large protein embedding language model like ESM2 [14], and *(iii)* initial within species cell annotations (from cell type assignments if available or obtained by running a clustering algorithm). The language model takes a sequence of amino acids and produces a protein representation vector (Fig. 1a). Given gene expression and protein embeddings, SATURN learns an interpretable feature space shared between multiple species. We refer to this space as a macrogene space and it represents a joint space composed of genes inferred to be functionally related based on the similarity of their protein embeddings. The importance of a gene to a macrogene is defined by a neural network weight – the stronger the importance, the higher the value of the weight that connects the gene to the macrogene.

**Figure 1.**
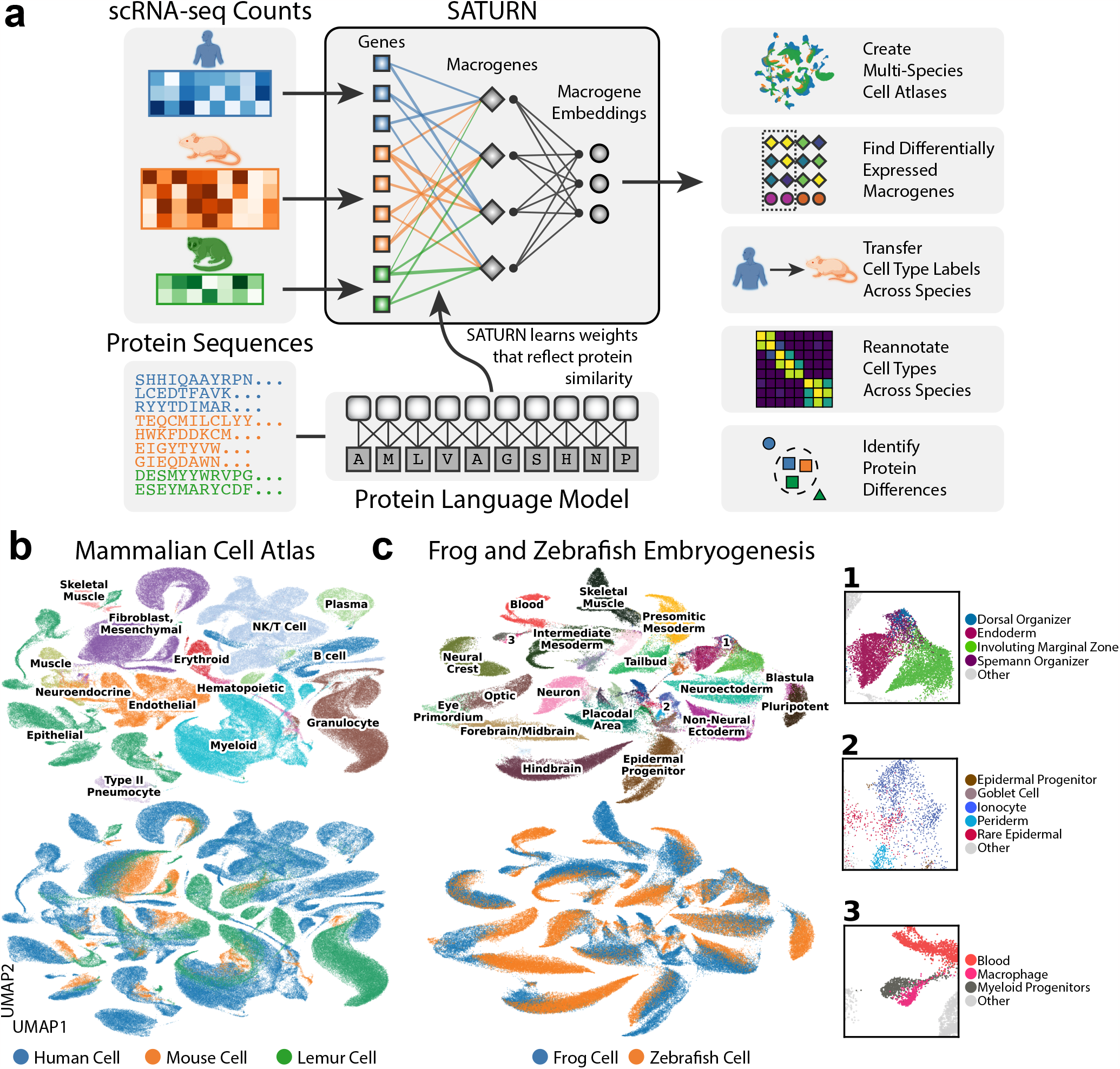
SATURN incorporates protein sequences and gene expression to embed single cells. **(a)** Overview of SATURN. SATURN takes as input scRNA-seq datasets generated from one or more species and the amino acid sequences of proteins present in these species. SATURN then maps each species’ genes to a joint feature space by learning “macrogenes”– groups of functionally related intra- and inter-species genes. Finally, in the shared macrogene space SATURN integrates datasets across species by learning a joint cell embedding space in which cell types conserved across species are aligned with each other. **(b)** UMAP visualization of a joint embedding space across three distinct species. We apply SATURN to integrate cell atlas datasets of 335, 000 cells from Tabula Sapiens (human), Tabula Microcebus (mouse lemur) and Tabula Muris (mouse), creating a mammalian cell atlas. Colors denote coarse-grained cell type annotations (top) and species annotations (bottom). Only cell types with more than 350 cells were included. **(c)** UMAP visualization of SATURN ‘s integration of datasets from frog (97, 000 cells) and zebrafish (63, 000 cells) embryogenesis. Colors denote different major cell types (top) and different species (bottom). In SATURN ‘s embedding space, cell types conserved across species align well (*e*.*g*., frog/zebrafish neural crest) while species-specific cell types form separate single species clusters (*e*.*g*., frog goblet cells). Cell types not directly mapped between both species share similar ontology, for example the zebrafish dorsal organizer and frog Spemann organizer (inset 1). Epidermal cell types including periderm, epidermal progenitor, and rare epidermal cell types are also aligned, as are specialized epithelial cells such as goblet cells and ionocytes (inset 2). Finally, myeloid cell types including macrophages and myeloid progenitors cluster together (inset 3).

Given the shared macrogene expression space across different species, SATURN then learns to represent cells across multiple species as nonlinear combinations of macrogenes. The neural network in SATURN is first pretrained with an autoencoder with zero inflated negative binomial (ZINB) loss, regularized to reconstruct protein embedding similarities using gene to macrogene weights (Methods). Using the pretrained network as initialization, SATURN then learns a mapping of all cells to the shared embedding space with a weakly supervised metric learning objective. This allows SATURN to calibrate distances in the embedding space to reflect cell label similarity. In particular, the objective function in SATURN consists of two main components: *(i)* forcing different cells within the same dataset far apart using weak supervision; and *(ii)* forcing similar cells across datasets close to each other in an unsupervised manner (Methods). This objective enables SATURN to integrate cells across different species, while preserving cell type information within each species’ dataset.

### SATURN creates multi-species cell atlases

We applied SATURN to integrate large-scale single cell atlas datasets generated from human (Tabula Sapiens), mouse lemur (Tabula Microcebus) and mouse (Tabula Muris), creating the mammalian cell atlas of 335, 000 cells (Fig. 1b, Supplementary Fig. 1a). We found that major cell types aligned well across three species such as T cells, B cells and muscle cells; and then analyzed the alignment on a per-tissue level. For example, in muscle, we found a small subcluster of cells labeled as mouse macrophages that grouped with human and lemur granulocytes while the rest of cells labeled as mouse macrophages aligned with human and lemur macrophages (Supplementary Fig. 2, Supplementary Fig. 3). To investigate whether this alignment is indeed correct, we checked the expression of known granulocyte marker *Cd55* [19] [20] and known macrophage marker *Cd74* [19] [20]. Interestingly, we found that this small subcluster labeled as mouse macrophage indeed expresses *Cd55* and does not express *Cd74*, indicating that this small cluster was wrongly annotated as macrophages while it should be annotated as granulocyte (Supplementary Fig. 3).

In spleen, SATURN separated out human naive B cells from human memory B cells, but aligned human memory B cells with cells annotated as B cells in mouse and lemur (Supplementary Fig. 4, Supplementary Fig. 5). To investigate whether this alignment is meaningful, we checked the marker genes and found that indeed mouse and lemur B cells express *Cd19*, a B cell marker known to be preferentially expressed in memory B cells, which was only weakly expressed in human naive B cells (Supplementary Fig. 5) [21]. This indicates that mouse and lemur B cells are correctly clustered with human memory B cells, which is additionally confirmed by strong expression of *Cd19*. Thus, SATURN can be used to obtain fine-grained level annotations when cell atlases have been annotated with different granularity levels. Additionally, we found that SATURN correctly identified cell types specific to a single species within the integrated datasets. For instance, in muscle tissue, SATURN separated human epithelial and mesothelial cells from all other cell types (Supplementary Fig. 2). These cell types are indeed absent in mouse and lemur datasets. In spleen, SATURN separated human erythrocytes (Supplementary Fig. 4).

We next applied SATURN to a multi species dataset of frog (97, 000 cells) and zebrafish (63, 000 cells) embryogenesis [17]. SATURN aligned evolutionary related cell types between these two remote species (Fig. 1c, Supplementary Fig. 1b). We further inspected small clusters that are aligned by SATURN, but their ground-truth cell type annotations differ. We find that these clusters indeed correspond to related cell types. For example, SATURN integrated zebrafish early stage macrophages and frog myeloid progenitors which can differentiate into macrophages. Terminal differentiation in both cell types involves activation of a number of conserved master regulatory genes, such as *Cybb, Cyba, Spib*, and *Cepba* [17]. These cell types are embedded close to blood cells which further demonstrates that local distances in SATURN ‘s embedding space are meaningful.

### SATURN performs differential expression on macrogenes

SATURN extends differential expression analysis to a multi species setting. Instead of performing differential expression analysis on individual genes, which is highly limited when datasets do not share genes, SATURN performs differential expression on macrogenes, which enables characterization of cell-type specific macrogenes across different datasets. To perform differential expression on macrogenes, SATURN first aggregates the contributions of individual genes to macrogenes using gene-macrogene neural network weights (Fig. 2a). The aggregated values can be seen as macrogene expression for each individual cell. Like in conventional differential expression analysis, SATURN then performs differential expression on cell clusters, such as those determined by cell type label. The difference compared to conventional differential expression is that in SATURN the statistical test is performed on the macrogenes. Finally, to interpret the biological meaning of a macrogene, SATURN considers genes with the highest weight to the macrogene. We note that mean expression of a gene does not affect its macrogene weight. In particular, in the frog and zebrafish embryogenesis datasets, the correlation between a gene’s expression and its maximum weight is 0.08 and 0.05 in the frog and zebrafish datasets, respectively.

**Figure 2.**
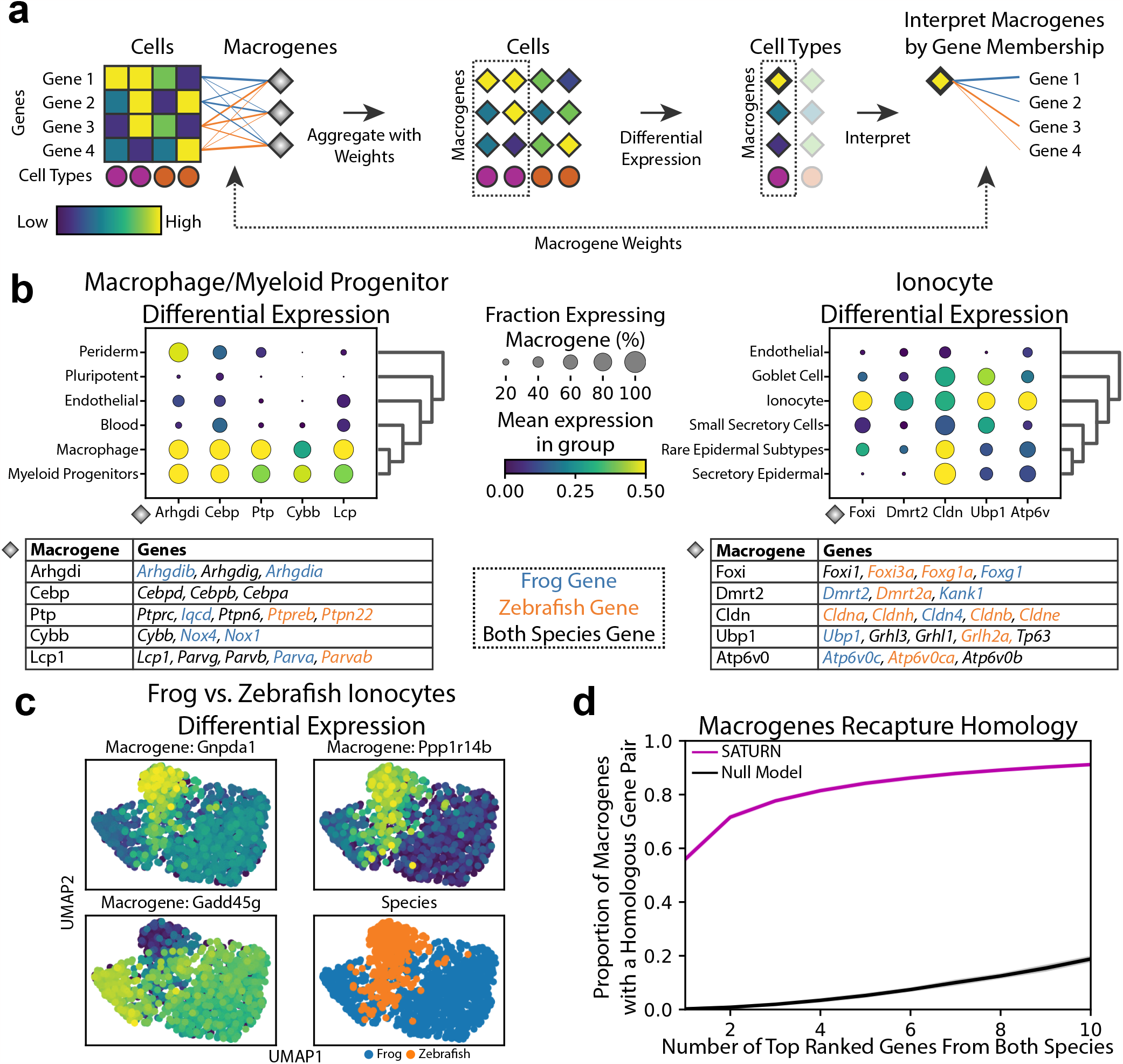
SATURN enables multi species differential expression analysis in the macrogene space. **(a)** Overview of SATURN ‘s differential expression analysis on macrogenes. Every gene is connected to a macrogene with a corresponding weight that represents the importance of that gene to the given macrogene. Thus, each cell has corresponding macrogene values calculated as the weighted and normalized sum of its gene expression values. Since SATURN operates in the macrogene space, differential expression for resulting cell clusters gives the set of differentially expressed macrogenes of a given cell type. Finally, the genes with the highest weights to a macrogene are used to determine the biological interpretation of that macrogene. **(b)** Differentially expressed macrogenes on frog and zebrafish embryogenesis datasets for **(left)** macrophage and myeloid progenitors and **(right)** ionocytes. Differential expression is performed by comparing these cell types with all other cell types. We show only cell types that are similar to target cell types determined as expressing a subset of the top differentially expressed macrogenes. We assign names to macrogenes based on the set of genes with the highest weight in the given macrogene. The tables show the top 5 differentially expressed macrogenes and the top weighted genes in each macrogene. Genes are shown in a black color if a gene is included in the top genes for both species in a given macrogene, and blue or orange if the gene is frog or zebrafish specific, respectively. **(c)** Macrogene differential expression can also be used to find species-level differences between cell types conserved across species. Example of differentially expressed macrogenes between frog and zebrafish ionocytes. **(d)** SATURN macrogenes contain a far higher proportion of homolog gene pairs than what is expected by chance, demonstrating that SATURN recaptures sequence based homology. Purple curve shows the proportion of SATURN macrogenes which contain, within their top ranked frog and top ranked zebrafish genes, at least one homolog gene pair, vs the top *N* number of genes. Homology is determined according to BLASTP results. The black curve shows the proportion obtained by a null model in which the same number of genes are randomly selected without replacement from both species. Error bars are the standard error from 30 independent experiments and they are nearly zero so not visible.

By performing macrogene differential expression SATURN has two major advantages over existing integration methods. First, SATURN can identify differentially expressed genes that lack a one-to-one homolog. This is in contrast to existing methods that rely on one-to-one homologs and therefore ignore unmapped genes. Second, differentially expressed macrogenes provide natural gene modules that aid in interpretation, since they rely on groups of related genes instead of individual genes. This can lead to identification of shared gene programs across species.

We conduct macrogene differential expression analysis in frog and zebrafish embryogenesis datasets. We demonstrate examples for the macrophage/myeloid progenitor cluster (Fig. 2b) and for the ionocytes cluster (Fig. 2b). In particular, we show the top five differentially expressed macrogenes and their corresponding highly weighted genes that characterize them, and we name each macrogene according to the gene with the highest weight to that macrogene. We focus on genes with known annotations. Gene to macrogene weights are listed in Supplementary Table 1.

For both macrophage/myeloid progenitors and ionocyte cell types we find that highly expressed macrogenes indeed capture groups of related genes that are known to have the function associated with these cell types. In particular, for macrophage/myeloid progenitors, the top differentially expressed macrogenes include *Arhgdi, Cebp, Ptp, Cybb* and *Lcp1* (Fig. 2b). All these macrogenes contain genes associated with functions in blood cells. For example, the *Arhgdi* macrogene contains frog and zebrafish homologs of *Arhgdig*, as well as frog specific paralogs such as *Arhgdib* and *Arhgdia*, which are involved in Rho protein signal transduction and RacGTPase binding activity [22, 23]. RhoGTPases play an important role in hematopoietic stem cell functions [24]. Similarly, the *Cebp* macrogene contains frog and zebrafish homologs of *Cebpd, Cebpb* and *Cebpa. Cebpa* is associated with zebrafish hemopoiesis and *Cebpb* is known to be expressed in zebrafish macrophages [22, 23].

For ionocytes, SATURN ranks *Foxi, Dmrt2, Cldn, Ubp1* and *Atp6v0* as top five differentially expressed macrogenes (Fig. 2b). Indeed, we find that all these macrogenes contain genes that are known to be associated with ionocytes. *Foxi* consists of Fox transcription factors that are known ionocyte markers [25]. The *Dmrt2* macrogene contains *Dmrt2* and *Dmrt2a. Dmrt2* is a known ionocyte marker in human pulmonary ionocytes [26]. The *Cldn* macrogene contains various claudins, which are found in gill ionocytes of teleost fish like zebrafish [27]. SATURN’s identification of a claudin marker macrogene for ionocytes is notable because the set of genes that can be mapped as one-to-one homologs does not contain any of these genes. Additionally, claudins that can be mapped as one-to-one homologs (*Cldn1, Cldn12, Cldn18, Cldn19* and *Cldn2*) are not differentially expressed within the top 200 differentially expressed ionocyte genes in the individual datasets, nor in the shared one-to-one homolog space.

Moreover, macrogene differential expression can also be used to find species-level differences between cell types conserved across species. For example, when comparing zebrafish and frog ionocytes a macrogene represented by *Gnpda1, Apip, Paics* and a macrogene represented by *Ppp1r14b* and *Fosab* are specific to zebrafish, while a macrogene represented by *Gadd45g, Aen*, and *Msgn1* is highly expressed in frog ionocytes but not in zebrafish (Fig. 2c). To analyze the proportion of single species versus species shared macrogenes, we found the top 20 differentially expressed macrogenes and then calculated the proportion of macrogenes that only had weights above 0.5 to one species’ genes. Across all cell types, 35% of macrogenes were represented by a single species’ genes.

### Macrogenes capture homology

We find that macrogenes generated by SATURN recapture sequence-based gene homologs. In particular, we computed the proportion of macrogenes with a homologous gene pair between zebrafish and frog among their top ranked genes. To assess gene homology, we use BLASTP, which determines gene homologs based on protein amino acid sequence similarity [28]. We find that even with only the top ranked genes of each species, 56% macrogenes in SATURN recapture gene homology information, while by considering ten top ranked genes from each species, 91.2% of macrogenes recapture gene homology information (Fig. 2d). In comparison, random assignment of genes to macrogenes results in homologous pairs in only 0.25% of macrogenes when considering two top ranked genes and in only 18.8% macrogenes when considering ten top ranked genes. Altogether, these results not only indicate that macrogenes in SATURN recapture homology information, but that they can also be used to reveal functional similarities between genes even when these genes are not considered as homologs by sequence based similarity tools such as BLASTP. To further demonstrate that macrogenes capture functional similarities of genes, we performed Gene Ontology (GO) [29] analysis between the human and mouse genes in the mammalian cell atlas datasets. The analysis revealed significantly enriched GO terms within the gene sets of the same macrogene (Supplementary Note 5).

### SATURN outperforms other methods by a large margin

We quantitatively assess the performance of SATURN on the alignment of frog and zebrafish embryogenesis datasets. We evaluate performance by measuring how well labels can be transferred from zebrafish to frog. In particular, we first integrate the datasets using SATURN and then use the cell-type annotations of cells from a reference species, zebrafish, to train a logistic classifier to predict cell types [30] (Supplementary Note 3). The classifier’s performance is then tested on the embeddings of the query species, frog (Fig. 3a). Predictions are assessed as correct if they match the known frog cell type, based on a predetermined mapping of cell types between species (Supplementary Table 2). Since not all frog cells can be mapped to zebrafish cells, the maximum possible accuracy of such a model is 93%.

**Figure 3.**
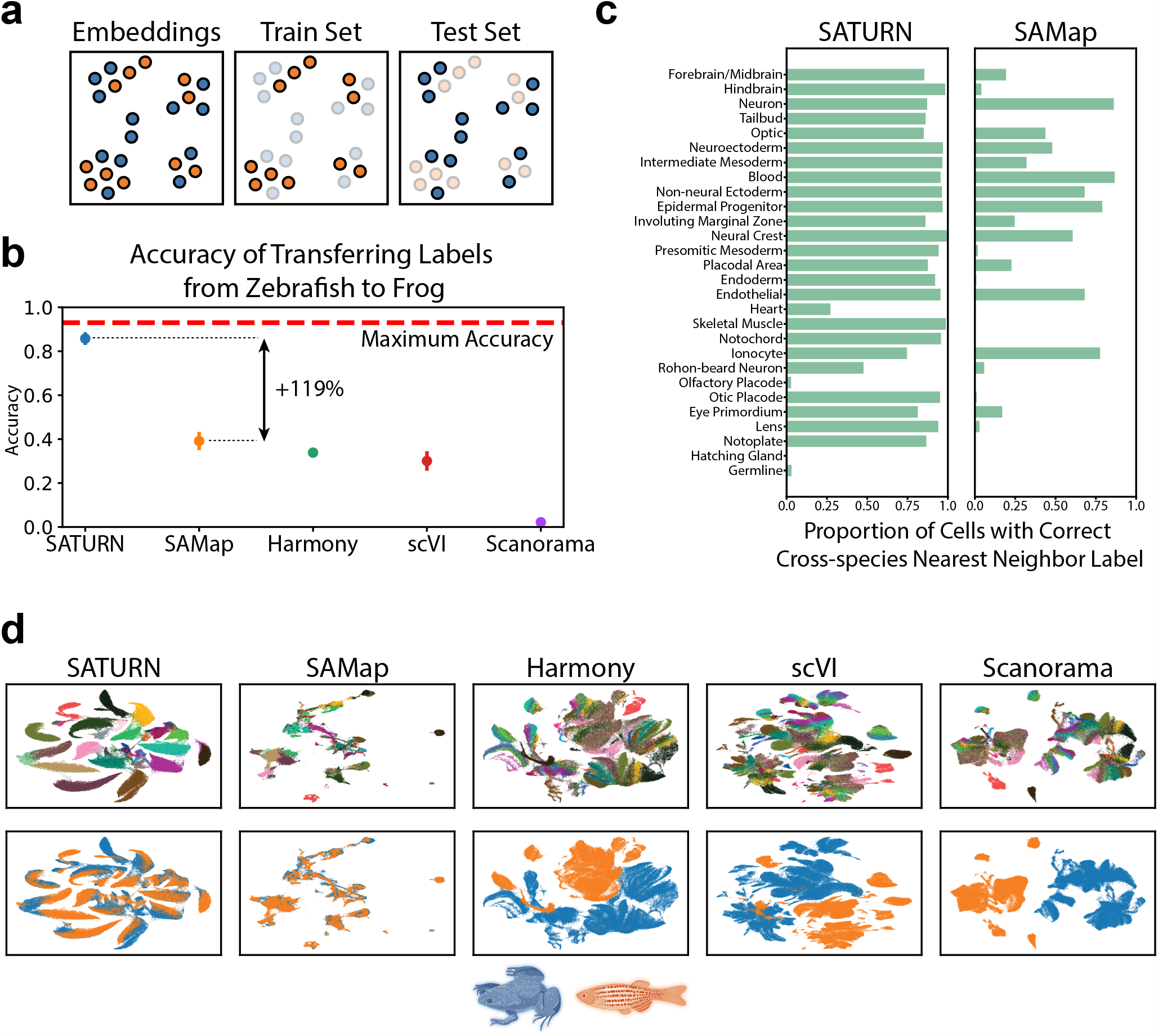
SATURN embeddings capture shared cell type identity in frog and zebrafish embryogenesis. **(a)** Explanation of how multi-species embeddings are scored. A joint embedding space, containing cells from multiple species, is split by species into a training set and a test set. A classification model to predict cell types is trained on a single species training set, and evaluated on the test set of another species. The maximum test set accuracy achievable will be lower than 100% if the test set species contains specific cell types that can not be predicted by a classifier trained on the training species. Blue color denotes frog, while orange denotes zebrafish. **(b)** Median performance of SATURN compared to alternative methods. The performance is evaluated using the prediction accuracy of a logistic classifier model trained to differentiate zebrafish cell types and tested on predicting the cell type annotations of frog cells. Higher values indicate better performance, and 0.93 is the maximum accuracy that can be reached by label transfer on this dataset. SAMap represents a version of the SAMap method in which cell-type annotations are used to integrate datasets. Vertical position of scatter plot points represents the median accuracy score across 30 runs for each method. Error bars represent standard error. For batch correction methods (Harmony, scVI and Scanorama), the input genes are selected as the one to one homologs determined by ENSEMBL. **(c)** SATURN produces more homogenous clusters than SAMap, and these clusters contain accurate multi species cell types. Bars represent the percentage of cells from zebrafish that are nearest neighbors of frog cells of the given cell type conserved across these two species. Cell types are ordered by frequency. **(d)** Comparison of UMAP visualizations of integrated frog and zebrafish embryogenesis datasets generated by SATURN and alternative methods. In SATURN ‘s embedding space different cell types naturally form clusters and cells from different species align well. On the other hand, alternative baselines either do not preserve cell type information (SAMap) or can not integrate two species (Harmony, scVI, Scanorama).

We compare the performance of SATURN to another single cell multi species integration method, SAMap [11], and unsupervised integration methods Harmony [6], scVI [8] and Scanorama [7]. SAMap is run in a weakly supervised mode in which cell neighborhoods are determined by cell type, which involves using the prior cell type label information within each species but not across species, which is the same setting we use for running SATURN. SAMap is initialized with a gene graph based on protein sequence similarity as determined by BLASTP. For the unsupervised methods, the input genes for each species are taken as the one-to-one homologs as determined by ENSEMBL [31]. We found that SATURN achieves 85.8% median accuracy in cell label transfer from zebrafish to frog, achieving remarkable 119% performance gain over the next best performing method, SAMap (Fig. 3b). We obtain similar performance gains when transferring labels from frog to zebrafish (Supplementary Fig. 6). Performance gains of SATURN are retained using other evaluation metrics, such as F1-score, precision and recall (Supplementary Fig. 7), as well as data integration metrics [32] (Supplementary Fig. 8). We additionally visualize embeddings obtained by using the dimensionality reduction techniques PCA and UMAP [33] on the one-to-one homolog expression space, demonstrating the gap between the species (Supplementary Fig. 9).

To test whether choice of protein language model for obtaining protein embeddings affects SATURN’s performance, we compared ESM2 embeddings [14] to ESM1b [12] and ProtXL [13]. The results show that SATURN is highly robust to the choice of protein language model (Supplementary Fig. 10), as well as to the number of macrogenes (Supplementary Fig. 11). SATURN also outperforms the best baseline approach on the mammalian cell atlas dataset (Supplementary Fig. 12).

We further compare SATURN ‘s ability to generate cell clusters that reflect conserved cell types across species, to the best baseline approach (SAMap). For each frog cell type we analyze its cross-species neighborhood by computing the cell type frequency of its nearest cross-species neighbors in the embedding space. We find that SATURN generates cell clusters that are both species heterogeneous and cell type homogeneous (Fig. 3c). For the most commonly occurring cell types, SATURN ‘s neighborhoods are consistently highly homogeneous. On the other hand, this is not the case for SAMap where the neighborhoods are typically cell type heterogeneous. For example, forebrain/midbrain, hindbrain, optic and eye primordium clusters are intermixed using SAMap but clearly distinguished using SATURN. SATURN aligns rare cell types such as notoplate, which only has 339 frog cells and 115 zebrafish cells. For a few very rare cell types, such as germline, which has only 33 frog cells and 53 zebrafish cells, SATURN and SAMap both fail to align. SATURN and SAMap fail to directly align additional rare cell types such as olfactory placode and hatching gland. However, SATURN aligns these cell types to functionally related cell types: 77% of olfactory placode cells are mapped to placodal area for SATURN (37% for SAMap) and 66% of hatching gland cells are mapped to another component of the EVL, the periderm, which is not case with SAMap (36% epidermal progenitor, 33% blastula).

We visually inspected low-dimensional embeddings produced by SATURN and other baselines by projecting them into a two-dimensional UMAP space [33]. We find that in SATURN ‘s embedding space different cell types form separate clusters while cell types conserved across species are mixed together (Fig. 3d, Supplementary Fig. 1b). On the other hand, existing methods are not able to produce biologically meaningful cell embeddings that reflect evolutionary signatures. In particular, Harmony, scVI and Scanorama fail to integrate datasets across remote species. While SAMap is able to integrate datasets across species, the cell type information in its embedding space is no longer preserved and different cell types intermingle.

### SATURN integrates five species from the Aqueous Humor Outflow cell atlas

SATURN scales to large datasets and it can handle multiple datasets at once. We apply SATURN to integrate five species Cell Atlas of Human Trabecular Meshwork and Aqueous Outflow Structures (AH atlas) [18]. The AH atlas contains 50, 000 cells from human, cynomolgus macaque, rhesus macaque, mouse and pig. SATURN jointly aligns different species in the embedding space, identifying many conserved cell types between these species (Fig. 4a, Supplementary Fig. 1c). SATURN embeddings suggest that cell types including melanocytes, macrophages, and ciliary muscle align in all species, as do cell types that are present only in a subset of species like fibroblasts and collector channel.

**Figure 4.**
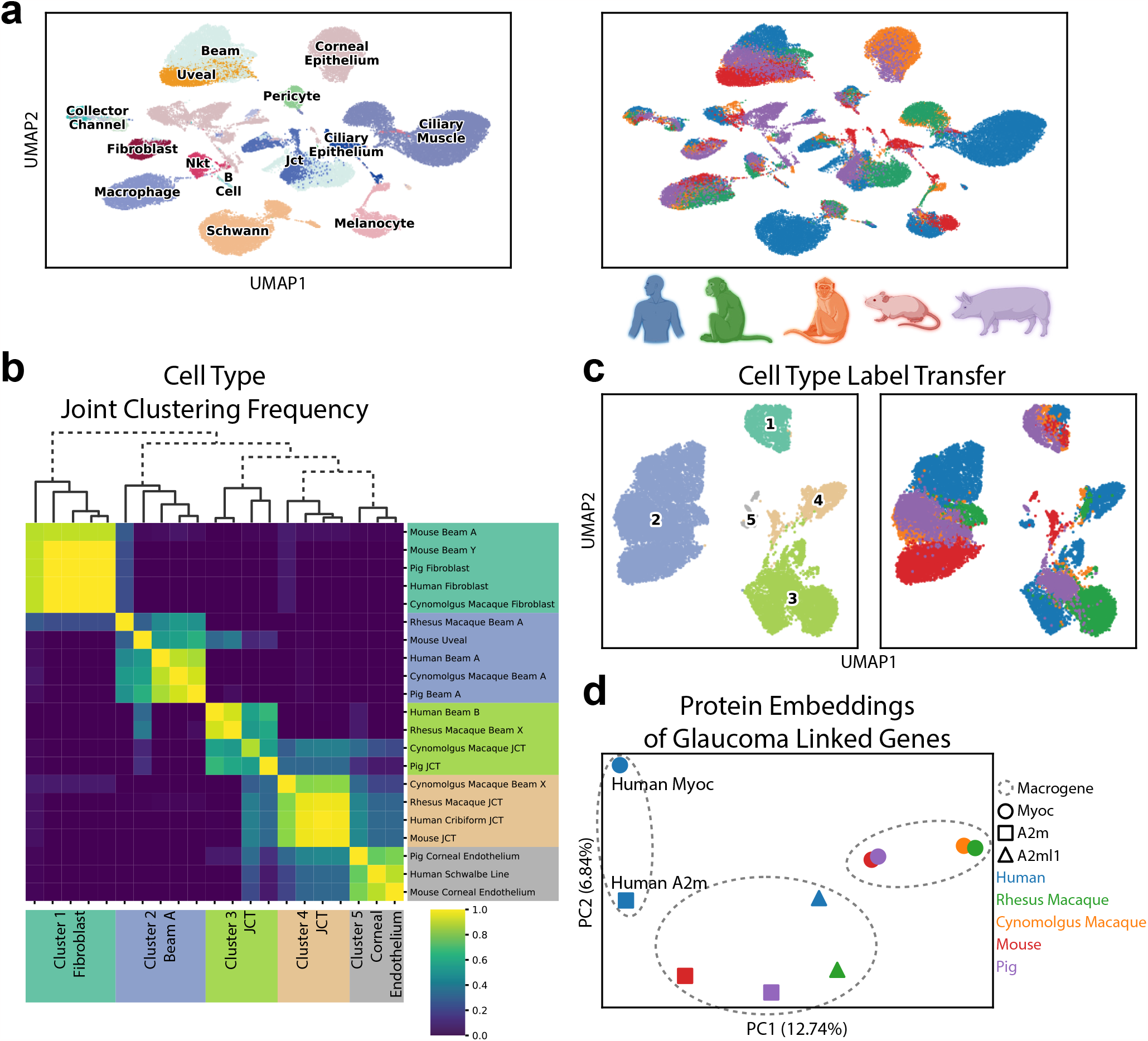
SATURN discovers new cell types and facilitates the analysis of protein embeddings for the Aqueous Humor Outflow cell atlas. **(a)** SATURN successfully aligns 50, 000 cells from the Aqueous Humor (AH) Outflow cell atlas consisting of five species: human, cynomolgus macaque, rhesus macaque, mouse, and pig. UMAP visualization of SATURN ‘s embeddings where colors denote cell types (left) and species (right). **(b, c)** We apply SATURN to regroup cell types in a multi-species context. By clustering SATURN ‘s embeddings, we find five broad cell types. **(b)** Heatmap and dendrogram of reannotated cell types using SATURN. Labels on the right side show original cell type annotations while on the bottom we show reannotations obtained using SATURN. These clusters include cell types originally labeled as fibroblast and beam A/Y cells (cluster 1), beam A and uveal cells (cluster 2), JCT and beam cells (cluster 3 and cluster 4), and corneal endothelium cells (cluster 5). Across 30 independent experiments, we regroup cluster 1 as fibroblast cells, cluster 2 as Beam A cells, clusters 3 and 4 as JCT cells, and cluster 5 as corneal endothelium cells. We specifically reannotate mouse beam A and beam Y cells, which have high expression of fibroblast markers such as *Pi16, FBn1*, and *Mfap5* as originally noted [18]. We additionally regroup human beam B cells, which were not found in other species, as JCT cells. Finally, we map beam X cells, which were unique to Rhesus and Cynomolgus macaque, to two JCT clusters. **(c)** UMAP visualizations of reannoted cell types. Cells are colored according to annotations inferred by SATURN (left) and species information (right). **(d)** SATURN facilitates the analysis of protein embeddings by creation of multi-species macrogenes. Human *Myoc* has highest weight to a different macrogene than the other four species’ *Myoc* variants. The human gene *A2m* also has highest weight to the human *Myoc* macrogene. We can investigate this discrepancy by visualizing the protein embeddings of *Myoc* and *A2m* from all five species using PCA. This analysis offers potential to point to similar function in *A2m* as *Myoc* which would otherwise not be identified by sequence based homology, as well as potential differences in human *Myoc* and *Myoc* in the other four species.

SATURN can be used to reannotate cell types and correct for incomplete annotations by aligning datasets across multiple species. To demonstrate that, we use SATURN to regroup cell types from the original AH atlas in a multi-species context. We focus on beam cells (beam A/B/X/Y), fibroblasts, JCT (juxtacanalicular tissue) cells, and corneal endothelium cells, due to their differential conservation across the five species profiled in the atlas.

Among these 21 cell types, SATURN found 5 broad clusters. The first cluster contains mouse beam cells and fibroblasts from pig, human and cynomolgus macaque, which we relabel as fibroblasts. The reannotated mouse beam cells are indeed characterized as having high expression of fibroblast marker genes (Supplementary Fig. 13, Supplementary Table 3). The second cluster contains beam A cells from pig, human, macaque and a mouse uveal cluster, which we reannotate as beam A cells. The third and fourth clusters contain beam X, beam B and JCT cells, which we reannotate as JCT cells, as beam X cells were only found in the two macaque species and beam B cells were only found in human. The fifth cluster contains the human Schwalbe line cells, and pig and mouse corneal endothelium cells. Within these new cell type groupings, we find differentially expressed macrogenes that recapture specific cell type marker genes (Supplementary Fig. 13, Supplementary Table 3).

### SATURN helps in finding homologous genes with potentially different functions across species

We investigate the macrogenes corresponding to glaucoma associated genes from each species in the AH atlas. While pig, mouse, cynomolgus and rhesus macaque *Myoc* gene are expectedly linked to the same macrogene, we find that the human *Myoc* gene is not linked to that macrogene. We next visualize protein embeddings of glaucoma associated genes and find that the human *Myoc* gene is embedded further away from the *Myoc* genes of the other species (Fig. 4d). Interestingly, the human *Myoc* gene has the highest weight to a macrogene containing human *A2m*, which is a non homologous gene that has also been associated with glaucoma [34], and a number of different non human species’ genes such as mouse *Folr1*, mouse *Fbln2*, mouse *Srgn*, and pig *SCP2d1. A2m* genes from non-human species have the highest weights to the same macrogene. This analysis demonstrates that protein embeddings in SATURN and their association to macrogenes can be used to search for sequence-based gene homologs with potentially different functions across species and that SATURN can facilitate the analysis of protein embeddings through the creation of macrogenes.

## Discussion

SATURN represents the first model that combines protein embeddings generated using large protein language models with gene expression from scRNA-seq datasets. By coupling protein embeddings with gene expression, SATURN learns universal cell embeddings that bridge differences between individual single-cell experiments even when they have different genes.

SATURN has a unique ability to map heterogeneous datasets to an interpretable space of macrogenes that can group together functionally related genes across species. In SATURN, every gene has a weight to a macrogene which defines an importance of that gene to the macrogene. This enables SATURN to perform differential expression in the macrogene space and identify gene programs shared across different datasets. However, explicitly associating each macrogene with an interpretable function is not always possible due to the varied definitions of biological function across different contexts and scales, coupled with insufficient existing gene annotations.

SATURN represents cells as nonlinear combinations of macrogenes. To integrate datasets, the objective function introduced in SATURN learns distance metrics from weakly supervised data which forces cells to cluster according to their cell types. SATURN allows integration of datasets generated across multiple different species. SATURN is a scalable approach which makes it applicable to large-scale cross-species cell atlas datasets. Our approach also has important implications for the creation of new multi-omic machine learning methods, including those that incorporate protein assay information (e.g. CITE-seq [35]), genotype, or those that assay a limited section of the transcriptome (e.g. MERFISH [36]). For example, to improve machine learning methods that incorporate protein assay information, proteins could be represented using protein embeddings, rather than as indices. Protein embeddings could also be modified or personalized using jointly measured genotype information. For integration of spatial datasets that profile only a subset of a transcriptome, SATURN does not require subsetting them to a set of common genes, which is required by current methods.

On the other hand, the limitation of SATURN is the requirement of a reference proteome which may be missing for some species of interest. Reference proteomes and genomes can underrepresent the genetic diversity of species, even for well studied species such as humans [37]. More-over, to generate the protein embeddings used by SATURN, we average over the embeddings produced for each gene’s available protein products, ignoring various RNA splicing dynamics that affect the final translational products of genes. SATURN also requires cell clusters as an input for each dataset. These cell clusters could be created at various resolutions, which could limit the transferability of labels. Finally, smaller cell clusters, such as the germline cells in frog and zebrafish embryogenesis, are difficult to faithfully integrate.

SATURN generates cell embeddings that can be used for many downstream tasks. These tasks include but are not limited to dataset integration, discovery of conserved and species-specific cell types, differential macrogene expression analysis, cell type reannotation, signature set enrichment, gene module determination [38], or trajectory inference [39]. As single cell transcriptomics is applied to an increasing number of species, we expect SATURN will be an important tool for comprehending conservation and diversification of cell types across species and revealing fundamental evolutionary processes.

## Methods

### Overview of SATURN

SATURN takes multiple annotated single cell RNA expression count datasets generated from *S* species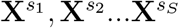 where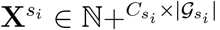 where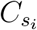 is the number of cells in species *s*_*i*_ and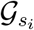 is the set of genes in species *s*_*i*_. The initial cell annotations can be obtained either from cell type assignments if available or obtained by running a clustering algorithm. In all experiments in the paper, we run SATURN with initial cell type assignments within the individual species but never matched across species. In addition to count matrices and cell type labels, SATURN also takes as input *p*-dimensional protein embeddings **P** ∈ ℝ^|*𝒢*|*×p*^ generated from large protein language models where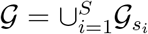.

SATURN maps multi-species expression data to a joint low dimensional macrogene expression space by learning a set of macrogenes *ℳ* with weights **W** ∈ ℝ+^|*𝒢*|*×ℳ*^ where **W**_*g,m*_ ∈ ℝ+ is a weight from a macrogene *m ∈ ℳ* to a gene *g ∈ 𝒢*. SATURN generates final *k*-dimensional latent cell embeddings by combining macrogenes using an encoder neural network *f* : ℝ^|*ℳ*|^ *→* ℝ^*k*^. SATURN consists of two main steps: (i) pretraining using an autoencoder, and (ii) fine-tuning using metric learning approach. Both steps are performed jointly on the datasets from all species.

### Macrogenes initialization

SATURN initializes macrogenes by soft-clustering protein embeddings. In particular, SATURN first clusters protein embeddings using the K-Means algorithm [40]. Given a matrix that stores protein embeddings for all genes **P** *∈* ℝ^|*𝒢*|*×p*^, SATURN applies K-Means to learn a set of centroids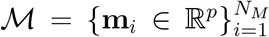 where *N*_*M*_ defines the number of centroids/macrogenes. K-means minimizes the within-cluster sum of squares:

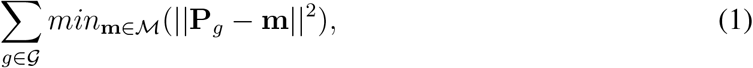

where **P**_*g*_ denotes a row protein embedding vector of matrix **P**. Here, each centroid **m** represents a different macrogene. SATURN then defines an initial set of weights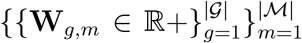 from each gene *g* to each macrogene *m* as:

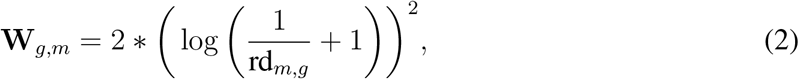

where rd_*m,g*_ : ℕ *→* ℕ represents the ranked euclidean distance from gene *g* to a macrogene *m* and rd_*m,g*_ = 1 for the nearest gene to a macrogene. This initialization function is arbitrarily chosen so that genes have the highest weights to the macrogenes they are closest to. Gene to macrogene weights are strictly positive, differentiable and updated during pretraining. We also explore different weight initialization strategies and show robustness of SATURN to different initialization functions (Supplementary Fig. 14, Supplementary Note 6). We multiply by two so that the highest weights are close to 1.

### Pretraining with an autoencoder

Following macrogene initialization, SATURN pretrains a network using an autoencoder with zero inflated negative binomial (ZINB) loss [8]. The autoencoder is composed of encoder and decoder modules. The encoder module first aggregates expression values using macrogene weights. In particular, for a cell *c* from species *s* with count values 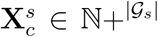, genes *g* ∈ *𝒢*_*s*_ and macrogenes *m* ∈ ℳ, SATURN defines macrogene expression values **e**_*c*_ ∈ ℝ+^|*ℳ*|^ as follows:

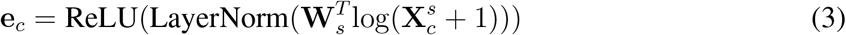

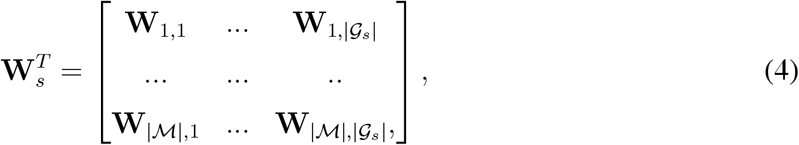

where ReLU denotes the rectified linear unit used as the activation function and defined as *ReLU* (*·*) = *max*(0, *·*). Macrogene expression values are always positive to ensure that each gene positively contributes to a macrogene or does not contribute at all. LayerNorm is layer normalization [41] defined as:

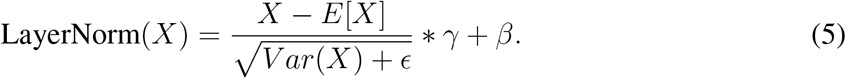

The encoder module *f* consists of two fully connected neural network layers with ReLU activation, layer normalization and dropout, and takes as an input **e**_*c*_ ∈ ℝ+ and outputs a low dimensional embedding **z**_*c*_ *∈* ℝ^*k*^ :

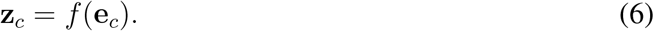

The decoding module outputs three distinct heads, parameterizing |*𝒢*| zero inflated negative binomial distributions with ***µ***_*c*_ *∈* ℝ+^|*𝒢*|^, **O**_*c*_ *∈* ℝ^|*𝒢*|^, ***θ*** *∈* ℝ+^|*𝒢*|^.

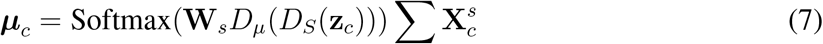

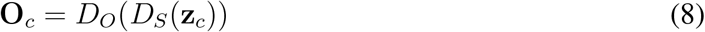

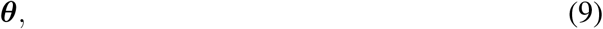

where *D*_*S*_, *D*_*µ*_ and *D*_*O*_ represent fully connected neural network layers. *D*_*S*_ and *D*_*µ*_ have ReLU activation, dropout and layer normalization. ***θ*** is a differentiable parameter of the model. SATURN provides the ability to concatenate a one hot representation of the species *s* to the embedding **z**_*c*_ in Eq. (6) during pretraining of the autoencoder. However we find that this does not improve the performance and set the species conditional variable to a constant value in all experiments (Supplementary Fig. 15). That including the species as a conditional variable does not improve performance may be of consideration for the development of other auto encoder based methods for single cell expression data. However, while performance was not helped in this case, for other settings, or datasets, a conditional variable autoencoder might be the correct choice, and we include the ability to pretrain with a CVAE in the SATURN codebase.

The autoencoder reconstruction loss *L*_*rc*_, is calculated as the negative log likelihood of a zero inflated negative binomial distribution [8] parameterized as:

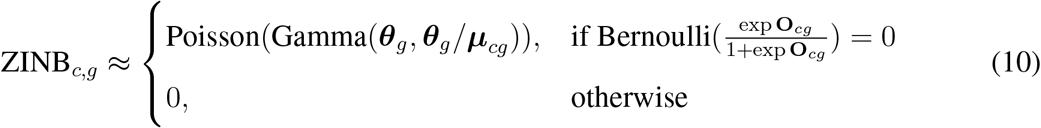

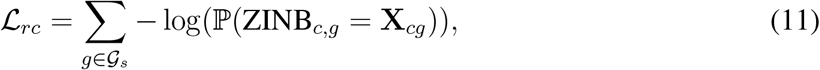

where ℙ denotes probability. To ensure that gene to macrogene weights reflect similarity in protein embedding space, we add an additional loss term *L*_*s*_ defined as follows:

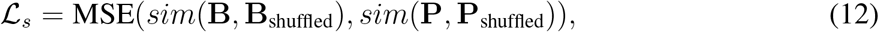

where **B** = *Q*(**W**) and *Q* : ℕ+^|*ℳ*|^ *→* ℕ^*n*^ is a fully connected neural network layer with ReLU activation, layer normalization and dropout, that encodes macrogene weights. MSE denotes mean squared error and *sim* is the cosine similarity. The encoded macrogene weights **B** *∈* ℝ^|*𝒢*|*×n*^ and protein embeddings **P** are jointly shuffled row-wise (gene-wise).

The final pretraining loss *ℒ*_*p*_ that SATURN optimizes is defined as:

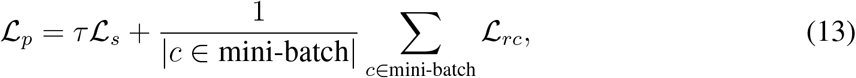

where *τ* is a regularization parameter and it is set to 1 in all experiments and mini-batch is a training mini-batch.

### Metric learning across species

To automatically learn a distance metric across species, SATURN fine-tunes pretrained cell embeddings with a weakly supervised metric learning objective. In particular, SATURN relies on the triplet margin loss function:

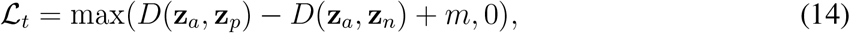

where *D* is a cosine distance, *a, p* and *n* denote an anchor cell, a positive cell and a negative cell, respectively, and the margin *m* is a tunable hyperparameter that we set to 0.2 in all experiments. Triplets are mined using semi-hard online mining in a weakly supervised fashion. To mine triplets, SATURN iterates over the species specific cell-type annotations, but no cross-species annotations are ever used. These within species annotations can be predetermined or generated in an unsupervised manner with clustering techniques like Leiden clustering [42]. For each annotation, SATURN selects all cells with that annotation from the same species as candidate anchor cells. Then for each anchor cell, SATURN selects candidate positive cells as mutual 1-nearest neighbors measured using cosine distance in the embedding space. Here, mutual means that if cell *x* from species *s*_1_ selected as its cross-species nearest neighbor cell *y* from species *s*_2_, SATURN finds the nearest neighbor *x*^*′*^ of cell *y* in species *s*_1_. If cells *x* and *x*^*′*^ from species *s*_1_ have the same annotation, then positive pairs are generated. The anchor cells and positive cells are pooled, and then matched such that each anchor cell candidate has a corresponding randomly selected positive cell candidate from a different species. Finally, negative cells are randomly selected such that they have a different label than either the anchor label or the positive label. Triplets are semi-hard filtered such that:

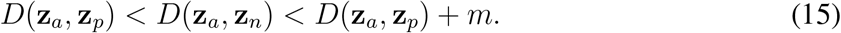

During the fine-tuning stage, macrogene weights are not updated.

### Generation of protein embeddings

Protein embeddings are generated by applying a pretrained protein embedding language model on each species’ reference proteome. Protein embeddings generated by the ESM2 model [14] were used for all experiments. The ESM2 protein embedding model accepts a sequence of amino acids as an input and outputs a *p* = 5120 dimensional vector representing the embedding of the protein. To obtain a protein embedding for a gene, the protein embeddings of all proteins available for the gene are averaged. Any protein embedding model, or any model that outputs numerical representations of genes, can be used as an input to SATURN (Supplementary Fig. 10).

### Differential macrogene expression

Differential expression on macrogene values is performed using a Wilcoxon rank sum test as implemented in SCANPY [43]. For a cell type annotation *t*, with cells *c ∈ t* (from any species), the test statistic *U*_*m*_ for macrogene *m* is calculated as:

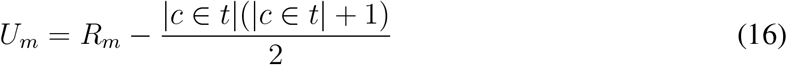

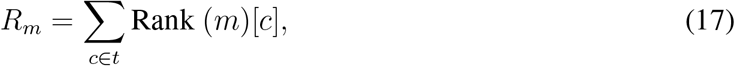

where *R*(*m*) is the rank sum of cells with label *t* for macrogene *m*.

### Determining gene homologs

BLASTP with default settings was applied to publicly available reference proteomes from ENSEMBL. BLASTP homologs results were used to find homolog gene pairs within the genes with highest weight to each macrogene (Fig. 2d). BLASTP results are also used for SAMap alignment (Fig. 3). The ENSEMBL homology API was queried to determine one-to-one gene homologs.

## Data availability

All analyzed datasets are publicly available. Tabula Sapiens is available at [CellXGene](https://cellxgene.cziscience.com/collections/e5f58829-1a66-40b5-a624-9046778e74f5). Tabula Microcebus is available at [FigShare](https://figshare.com/articles/dataset/Tabula_Microcebus_v1_0/14468196?file=31777475). Tabula Muris is available at [FigShare](https://figshare.com/articles/dataset/Single-cell_RNA-seq_data_from_microfluidic_emulsion_v2_/5968960/2). For embryogenesis datasets, frog is available with accession code GSE113074 and zebrafish is available in h5ad format at [KleinTools](https://kleintools.hms.harvard.edu/paper_websites/wagner_zebrafish_timecourse2018/WagnerScience2018.h5ad). The five species aqueous humor outflow pathway atlas datasets are available with accession code GSE146188.

## Code availability

SATURN was written in Python using the PyTorch library. The source code is available on Github at https://github.com/snap-stanford/saturn.

## Acknowledgements

We gratefully acknowledge the support of DARPA under Nos. HR00112190039 (TAMI), N660011924033 (MCS); ARO under Nos. W911NF-16-1-0342 (MURI), W911NF-16-1-0171 (DURIP); NSF under Nos. OAC-1835598 (CINES), OAC-1934578 (HDR), CCF-1918940 (Expeditions), NIH under No. 3U54HG010426-04S1 (HuBMAP), Stanford Data Science Initiative, Wu Tsai Neurosciences Institute, Amazon, Docomo, GSK, Hitachi, Intel, JPMorgan Chase, Juniper Networks, KDDI, NEC, and Toshiba. M.B. acknowledges the EPFL support. Created with BioRender.com.

## Author information

M.B., Y.RH. and J.L. conceived the study. Y.RS., M.B., Y.RH. and J.L. performed research, contributed new analytical tools, designed algorithmic framework, analyzed data and wrote the manuscript. Y.RS. performed experiments and developed the software. K.S. and Z.L. contributed to code.

