## Supplementary Materials for "Towards Universal Cell Embeddings: Integrating Single-cell RNA-seq Datasets across Species with SATURN"

This PDF file includes:

Supplementary Notes 1 to 6

Supplementary Figures 1 to 15

Supplementary Tables 1 to 3

Supplementary References

### Supplementary Note 1 Datasets and preprocessing

We downloaded publicly available count matrix files with cell type annotations (see data availability). For integrating Tabula Sapiens [1], Tabula Microcebus [2] and Tabula Muris [3] we filtered cell types to select cell types with more than 350 cells. Additionally, we filtered cells with fewer than 500 genes expressed and filtered genes expressed in fewer than 1000 cells. For frog and zebrafish embryogenesis, we filtered cells with fewer than 500 genes expressed, and filtered genes that were expressed in fewer than 10 cells. For the Aqueous Humor Outflow cell atlas no additional gene or cell filtering was done. We selected highly variable genes in each dataset using the Seurat v3 method [4]. We set only number of genes (Supplementary Note 4), while we keep all other parameters to their default values in scanpy package [5]. No additional data preprocessing was performed and the numerical inputs to SATURN are raw counts.

### Supplementary Note 2 Baseline methods

We compare SATURN to four existing single cell integration methods, SAMap, Harmony, scVI and Scanorama. SAMap is run in a semi-supervised mode in which cell neighborhoods are determined by cell types. Harmony [6], scVI [7], and Scanorama [8] are all run with default settings, using one-to-one homolog genes, and with the batch variable being species (frog or zebrafish). For scVI, Harmony and Scanorama, no additional highly variable gene selection was performed as the number of one-to-one homologs was low (7175). SAMap defaults to 3000 highly variable genes for each species, as determined by their SAM weights.

### Supplementary Note 3 Evaluation

There are a variety of different ways to assess the quality of a multi-species embedding. A multi-species embedding should encode cells that are the same cell type close together and cells from different cell types far apart. Cell types that are shared across species should have similar embeddings, and cell types that are unique to a species should not be falsely paired with other cell types.

We therefore assess the quality of multi-species embeddings for the goal of transferring labels from one species to another. Given a species $s^1$ with distinct cell types $T^1$, labels are transferred to a new species $s^2$ with cell types $T^2$ using a cell type classifier trained on the embeddings of cells from $s^1$. The simple classification model $C_{s^1}(\mathbf{z}_c) : \mathbb{R} \to T^1$ is trained on embeddings of one species $s^1$, and evaluated on embeddings of another species $s^2$. Predictions are classified as accurate based on a predetermined mapping of cell types $T^1 \to T^2$ between species (Supplementary Table 2).

$$C_{s^1} := \text{Logistic Regression Model}(\mathbf{z}_{c \in s^1}) \sim T^1_{c \in s^1} \tag{1}$$

$$\hat{T}^1_{c \in s^2} = C_{s^1}(\mathbf{z}_{c \in s^2}) \tag{2}$$

$$\text{Accuracy} = \frac{1}{|c \in s^2|} \sum_{c \in s^2} \mathbb{1}(\hat{T}^1_c \text{ maps to } T^2_c) \tag{3}$$

### Supplementary Note 4 Hyperparameters

**Hyperparameters.** In SATURN, we set the number of highly variable genes to $8000$. For integrating frog and zebrafish embryogenesis datasets and integrating the AH atlas, the number of macrogenes $|\mathcal{M}|$ is $2000$. For integrating tissue subsets of the mammalian atlas datasets, the number of macrogenes is $3000$. This dataset requires integration of fine-grained cell types from closely related species so we set the number of macrogenes to higher value. Intuitively, a higher number of macrogenes may help in finding finer-level differences between cell types, as an increased number of macrogenes will result in a more specific gene grouping. However, increasing the number of macrogenes past a certain point could reduce interpretability as the macrogenes may become too specific and consist of single genes. Since we are reducing the original high-dimensional gene space from all species in the macrogene space, we do not recommend using fewer than $1000$ macrogenes. The encoder embedding dimension, $k$ is set at $256$ for all experiments. The hidden dimension for all other layers used during pretraining is $256$. We use Adam optimizer with learning rate $0.0005$ during pretraining and $0.001$ during fine-tuning with metric learning.

To generate the coarse alignment of mammalian cell atlases in Fig. 1b, SATURN was run with $8000$ highly variable genes per species, $2000$ macrogenes, an embedding dimension $k$ of $256$ and a hidden dimension of $256$. An additional categorical covariate was added to the embedding dimension, representing the tissue of origin. All UMAP visualizations are generated using default values in scanpy package [5]. We generate UMAP embeddings with randomized plotting order in Supplementary Fig. 1.

### Supplementary Note 5 Gene Ontology enrichment analysis

Gene Ontology (GO) analysis could additionally confirm functionally meaningful groups of macrogenes. However, the challenge is that many species do not have well annotated GO terms and mapping GO terms across different species is non-trivial. Thus, we performed GO term enrichment analysis between human and mouse in the mammalian cell atlas, since human and mouse genes are best annotated in the GO. To create gene sets, for each macrogene we took the set of a given species' (either mouse or human) genes that had weights from a gene to macrogene above a cutoff of $0.5$. From these, to ensure gene sets had a sufficient size for enrichment analysis, we selected gene sets with $10$ or more genes, and ran GO enrichment analysis using the GOATOOLS Python package [9].

Using this approach, $88$ human gene sets and $79$ mouse gene sets were created. GO enrichment analysis on the human gene sets found an average of $2.05$ biological process (BP) terms, $1.35$ molecular function (MF) terms and $1.88$ cellular component (CC) terms that were enriched at a significant level (p=$0.05$, FDR BH corrected, default parameters) per human gene set. Enrichment analysis on the mouse gene sets found an average of $4.10$ BP terms, $2.38$ MF terms and $2.86$ CC significant terms per mouse gene set. In the null distribution of random assignment of genes, $0$ sets had significant terms of any kind. Moreover, we found $14$ macrogenes for which we could create gene sets for both human and mouse. In $11/14$ of these macrogenes we found at least one significantly enriched GO term in common between the mouse and human sets when performing string-based matching of terms.

### Supplementary Note 6 Macrogene initialization functions

**Default initialization.** By default, SATURN initializes macrogenes by soft-clustering protein embeddings. In particular, SATURN first clusters protein embeddings using the K-Means algorithm [10]. Given a matrix that stores protein embeddings for all genes $\mathbf{P} \in \mathbb{R}^{|\mathcal{G}| \times p}$, SATURN applies K-Means to learn a set of centroids $\mathcal{M} = \{\mathbf{m}_i \in \mathbb{R}^p\}_{i=1}^{N_M}$ where $N_M$ defines the number of centroids/macrogenes. K-means minimizes the within-cluster sum of squares:

$$\sum_{g \in \mathcal{G}} min_{\mathbf{m} \in \mathcal{M}}(||\mathbf{P}_g - \mathbf{m}||^2), \tag{4}$$

where $\mathbf{P}_g$ denotes a row protein embedding vector of matrix $\mathbf{P}$. Here, each centroid $\mathbf{m}$ represents a different macrogene. SATURN then defines an initial set of weights $\{\{\mathbf{W}_{g,m} \in \mathbb{R}+\}_{g=1}^{|\mathcal{G}|}\}_{m=1}^{|\mathcal{M}|}$ from each gene $g$ to each macrogene $m$ as:

$$\mathbf{W}_{g,m} = 2 * \left(\log\left(\frac{1}{\text{rd}_{m,g}} + 1\right)\right)^2, \tag{5}$$

where $\text{rd}_{m,g} : \mathbb{N} \to \mathbb{N}$ represents the ranked euclidean distance from gene $g$ to a macrogene $m$ and $\text{rd}_{m,g} = 1$ for the nearest gene to a macrogene. This initialization function is arbitrarily chosen so that genes have the highest weights to the macrogenes they are closest to. Gene to macrogene weights are strictly positive, differentiable and updated during pretraining. We multiply by two so that the highest weights are close to 1.

**Additional Functions.** We benchmark two additional initialization functions, a smoother function and an all-or-nothing "one-hot" function, which perform similarly (Supplementary Fig. 14).

For the more smoothed initialization function, the weights $\{\{\mathbf{W}_{g,m} \in \mathbb{R}+\}_{g=1}^{|\mathcal{G}|}\}_{m=1}^{|\mathcal{M}|}$ from each gene $g$ to each macrogene $m$ are set as:

$$\mathbf{W}_{g,m} = \frac{1}{\text{rd}_{m,g}} \tag{6}$$

For the one hot initialization function, the weights $\{\{\mathbf{W}_{g,m} \in \mathbb{R}+\}_{g=1}^{|\mathcal{G}|}\}_{m=1}^{|\mathcal{M}|}$ from each gene $g$ to each macrogene $m$ are set as:

$$\mathbf{W}_{g,m} = \mathbb{1}(\text{rd}_{m,g} = 1) \tag{7}$$

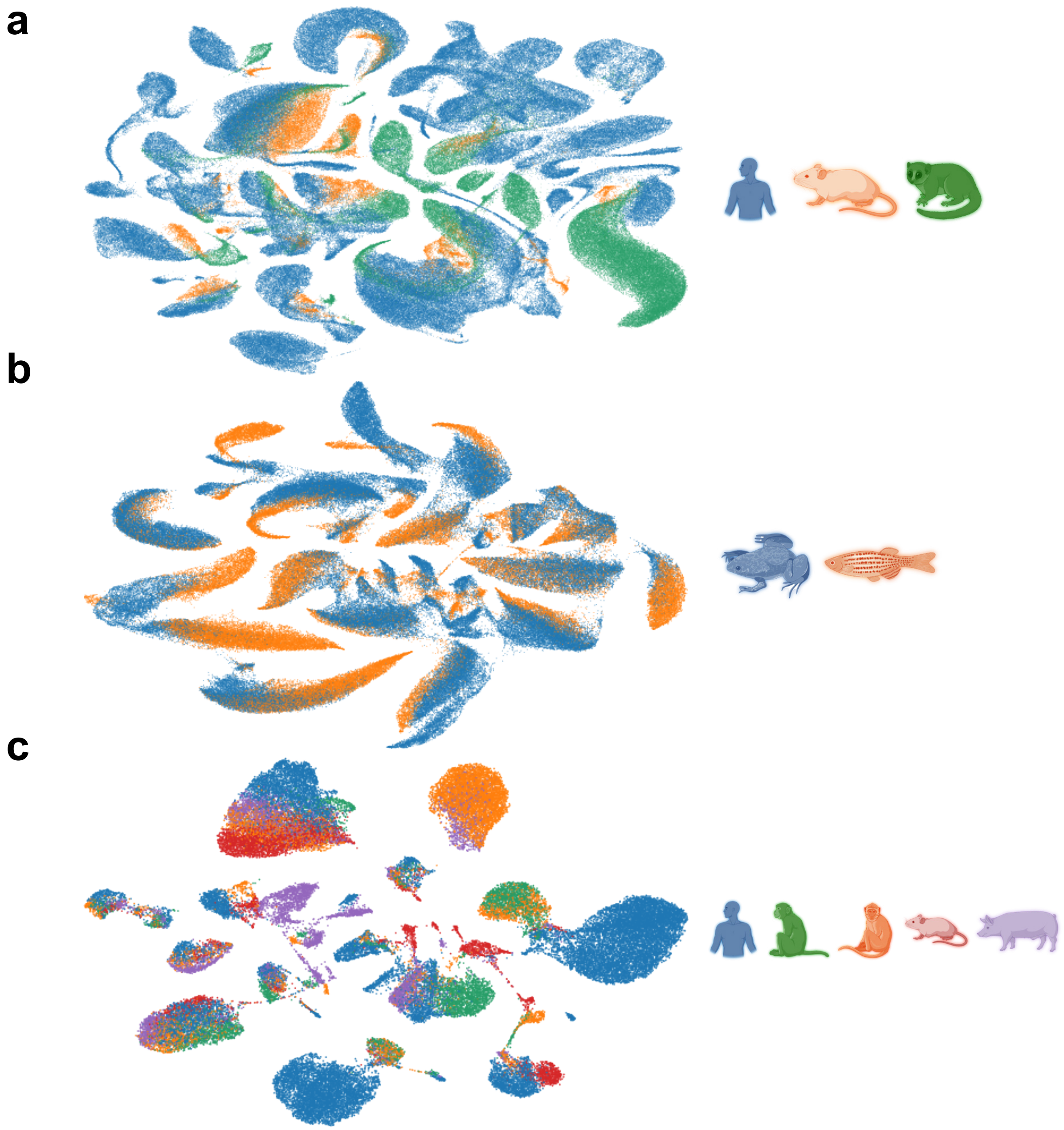


**Supplementary Figure 1: SATURN embeds multi species datasets.** UMAP embeddings of **(a)** mammalian cell atlas, **(b)** frog and zebrafish embryogenesis datasets and **(c)** Aqueous Humor Outflow cell atlas. UMAPs are generated using default parameters but plotting order is randomized.

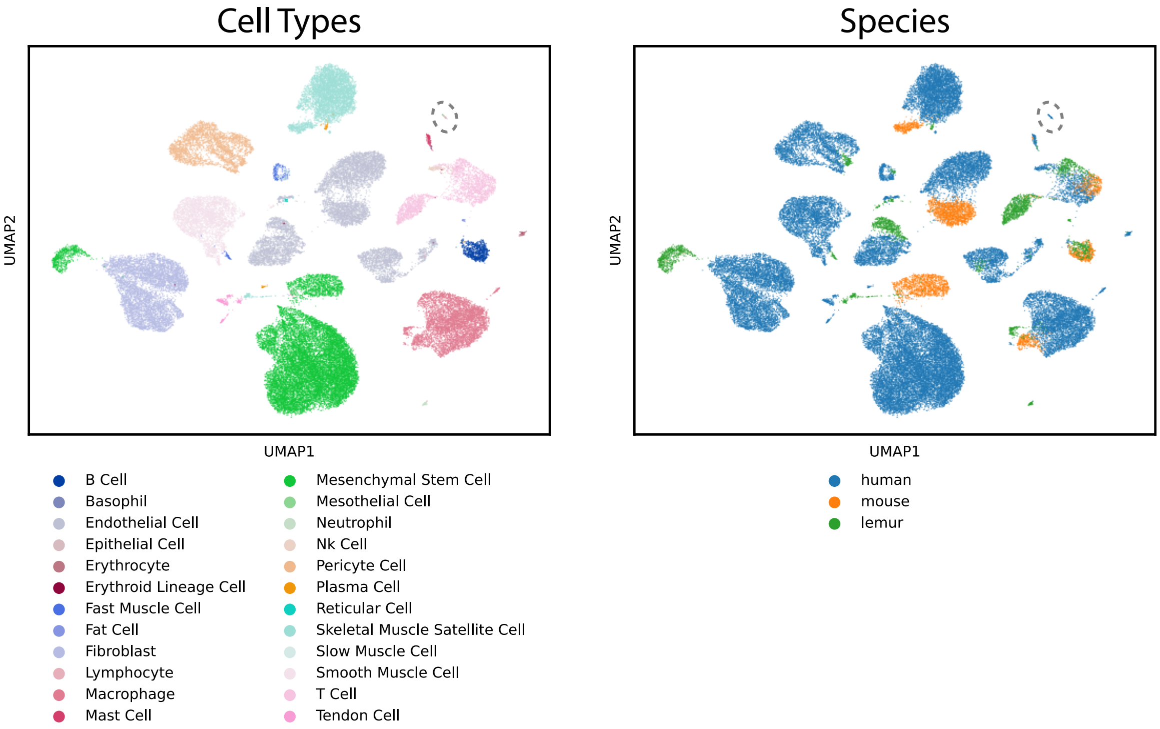


**Supplementary Figure 2: SATURN integrates muscle cell types across three mammalian species.** UMAP visualization of SATURN's embeddings obtained by integrating muscle tissues from human, mouse and lemur. Cells are colored based on broad-level cell types (left) and based on the species they come from (right). Epithelial and mesothelial cells types, which were only found within human, form a unique cluster (circled). To create each dataset, the larger Tabula datasets were subsetted. The human subset included cells labeled as muscle and vasculature. For mouse, limb muscle was chosen. For lemur, limb muscle and diaphragm were chosen. Human mesothelial cells belong to the tissue labeled muscle, and epithelial cells belong to the tissue labeled vasculature.

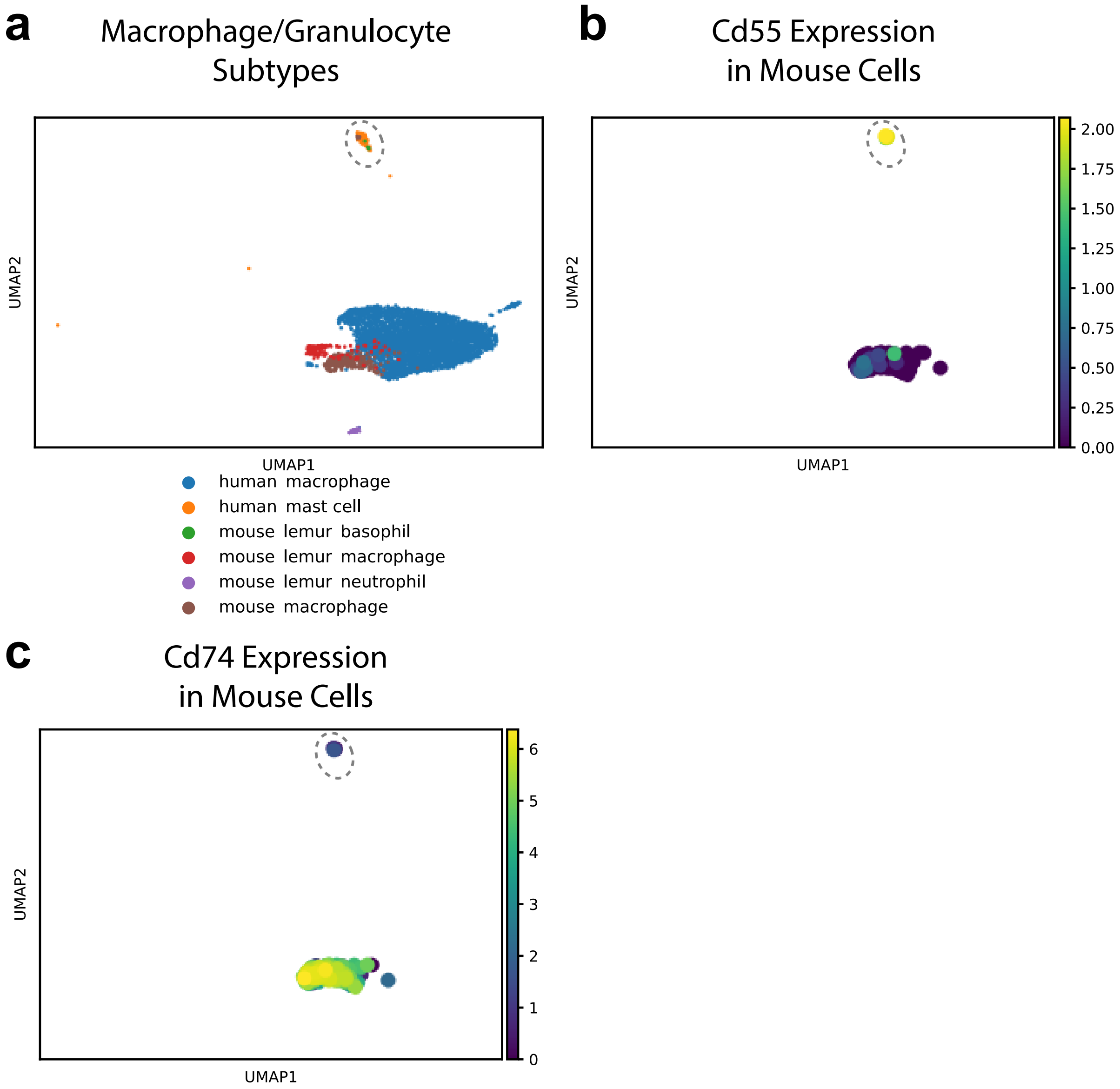


**Supplementary Figure 3: Reannotation of cells labeled as mouse macrophage in mouse muscle.** UMAP visualization of macrophage and granulocyte cell types obtained by integrating cells from human, mouse and lemur. **(a)** A small group of mouse macrophages cluster with granulocyte cell types from human (mast cells) and lemur (basophil) (circled), while other mouse macrophages cluster with human and lemur macrophages. These mouse cells **(b)** express *Cd55*, which has been shown to be preferentially expressed in granulocytes[11] [12], and **(c)** do not express *Cd74*, which has been shown to be preferentially expressed in macrophages and not expressed in granulocytes [11] [12]. **(b)(c)** are colored by log-normalized expression.

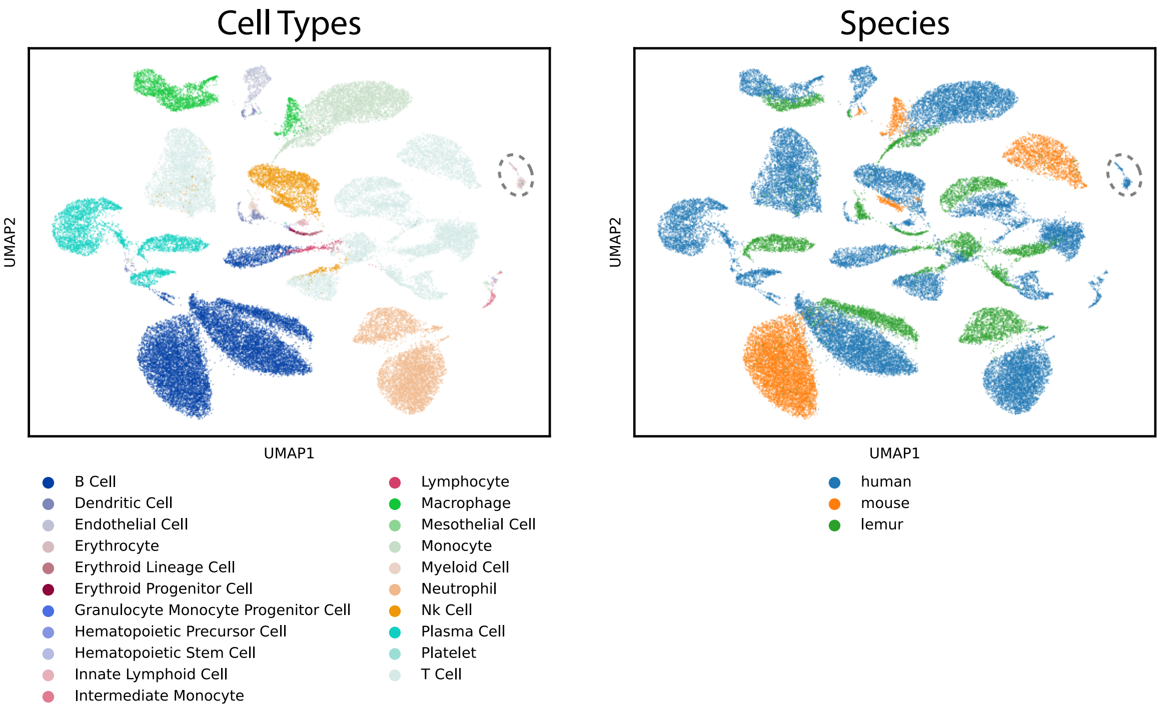


**Supplementary Figure 4: SATURN integrates spleen cell types across three mammalian species.** UMAP visualization of SATURN's embeddings obtained by integrating spleen tissues from the human, mouse and lemur. Cells are colored based on broad-level cell types (left) and based on the species they come from (right). Erythrocytes, which were only found within human, form a unique cluster (circled).

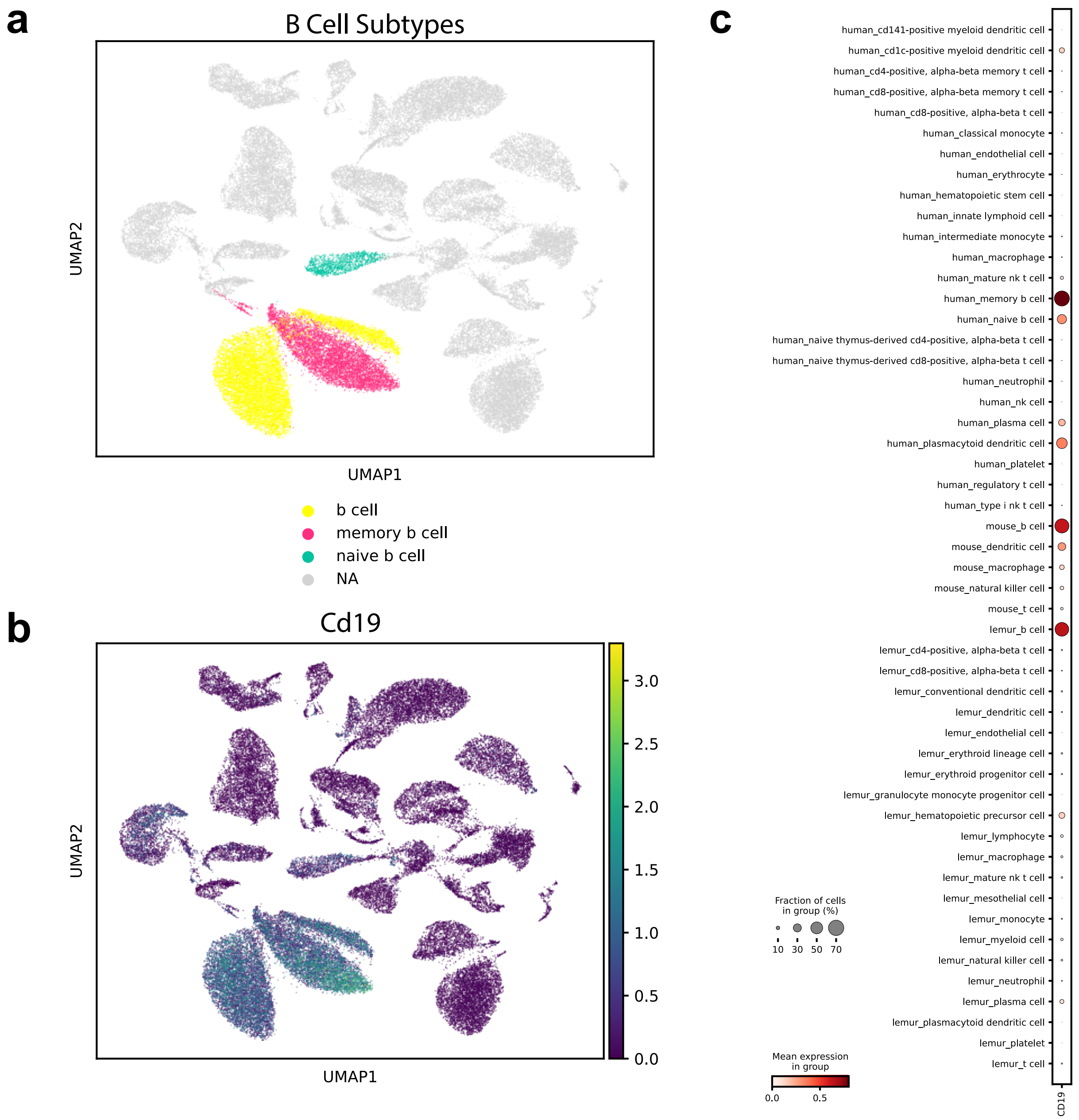


**Supplementary Figure 5: SATURN annotates B cells in mouse and lemur spleen on a fine-grained level.** **(a)** UMAP visualization of SATURN's embeddings obtained by integrating spleen cells from the human, mouse and lemur. B cells are shown in different colors based on ground-truth annotations while other cells are in grey. **(b)** UMAP visualization of expression of B cell marker *Cd19*. **(c)** Dotplot of *Cd19* expression vs species and cell type. *Cd19* is expressed in human memory B cells, mouse and lemur B cells, and only weakly in human naive B cells. This indicates that mouse and lemur B cells are correctly clustered with memory B cells.

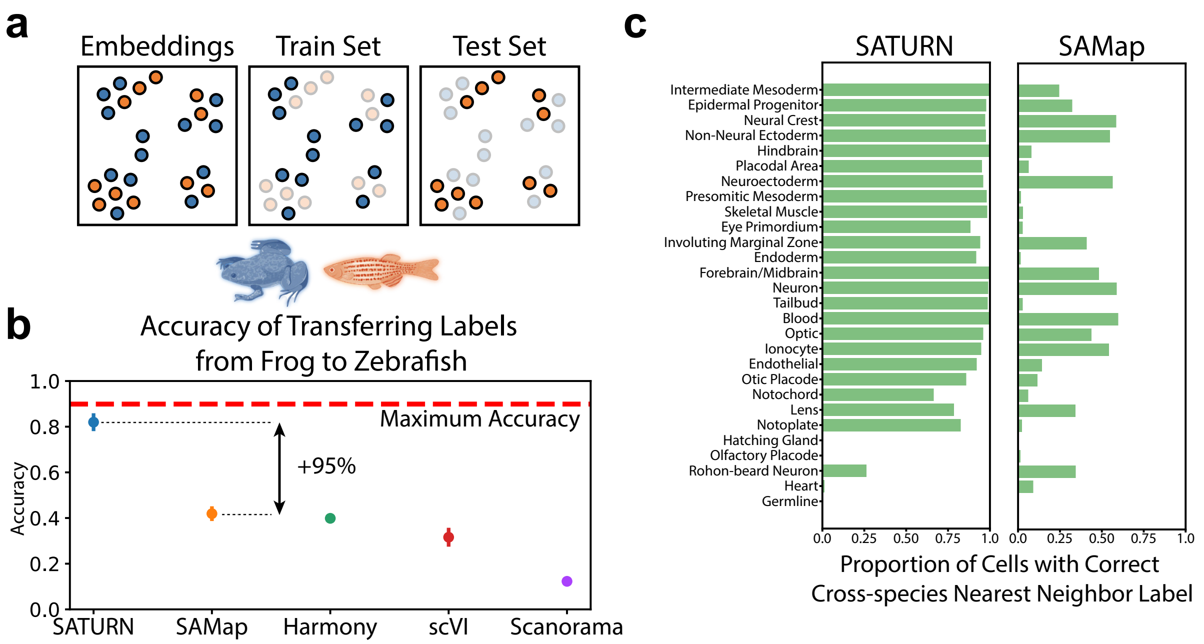


**Supplementary Figure 6: Label transfer from from frog to zebrafish embryogenesis datasets.** **(a)** Explanation of how multi-species embeddings are scored. A joint embedding space, containing cells from multiple species, is split by species into a training set and a test set. A classification model to predict cell types is trained on the frog training set cells, and evaluated on the zebrafish test set cells. The maximum test set accuracy achievable will be lower than 100% if the test set species contains specific cell types that can not be predicted by a classifier trained on the training species. Blue color denotes frog, while orange denotes zebrafish. **(b)** Median performance of SATURN compared to alternative methods. The performance is evaluated using the prediction accuracy of a logistic classifier model trained to differentiate frog cell types and tested on predicting the cell type annotations of zebrafish cells. Higher values indicate better performance, and 90% is the maximum accuracy that can be reached by label transfer on this dataset. SAMap represents a version of the SAMap method in which cell-type annotations are used to integrate datasets. Vertical position of scatter plot points represents the median accuracy score across 30 runs for each method. Error bars represent standard error. For batch correction methods (Harmony, scVI and Scanorama), the input genes are selected as the one to one homologs determined by ENSEMBL. **(c)** SATURN produces more homogeneous clusters than SAMap, and these clusters contain accurate multi species cell types. Bars represent the percentage of cells from frog that are nearest neighbors of zebrafish cells of the given cell type conserved across these two species. Cell types are ordered by frequency.

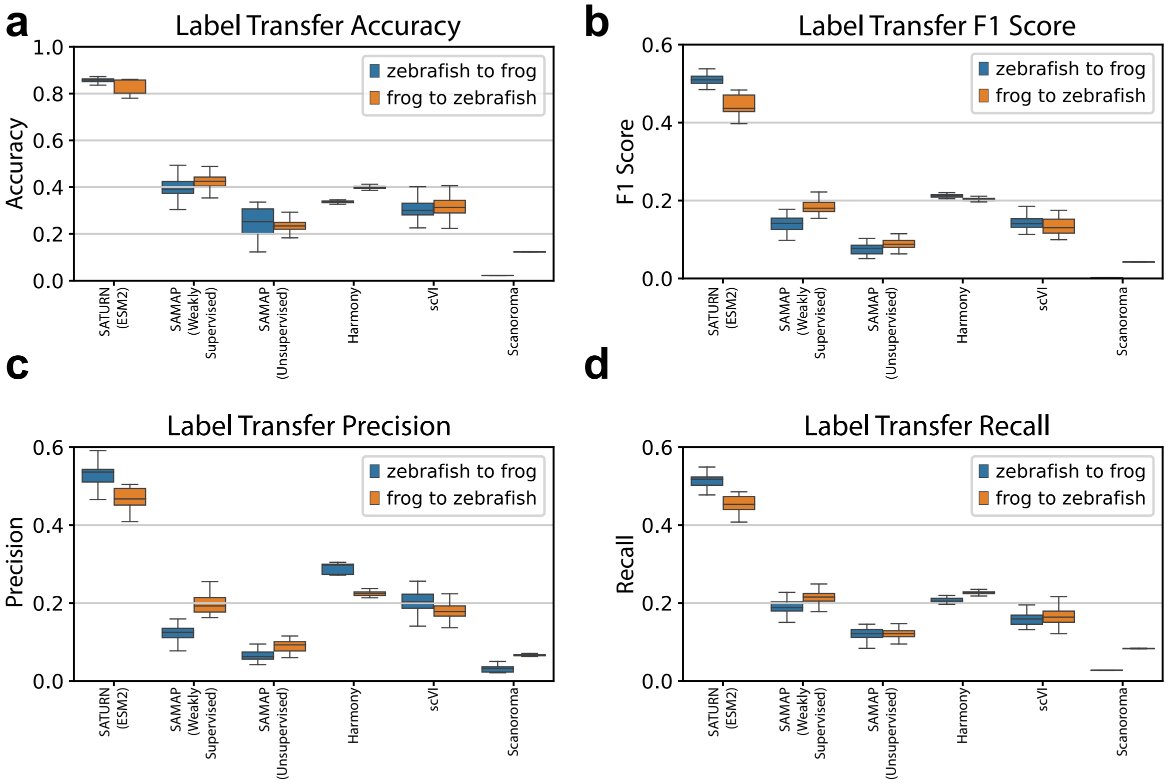


**Supplementary Figure 7: Performance comparison using different evaluation metrics.** Median performance of SATURN and baseline methods on label transfer between frog and zebrafish embryogenesis datasets evaluated using **(a)** accuracy, **(b)** macro-F1-score, **(c)** macro-precision, and **(d)** macro-recall. Blue boxplots show zebrafish to frog label transfer performance, while orange boxplots show frog to zebrafish label transfer performance. Distribution is estimated with $n = 30$ runs of each method.

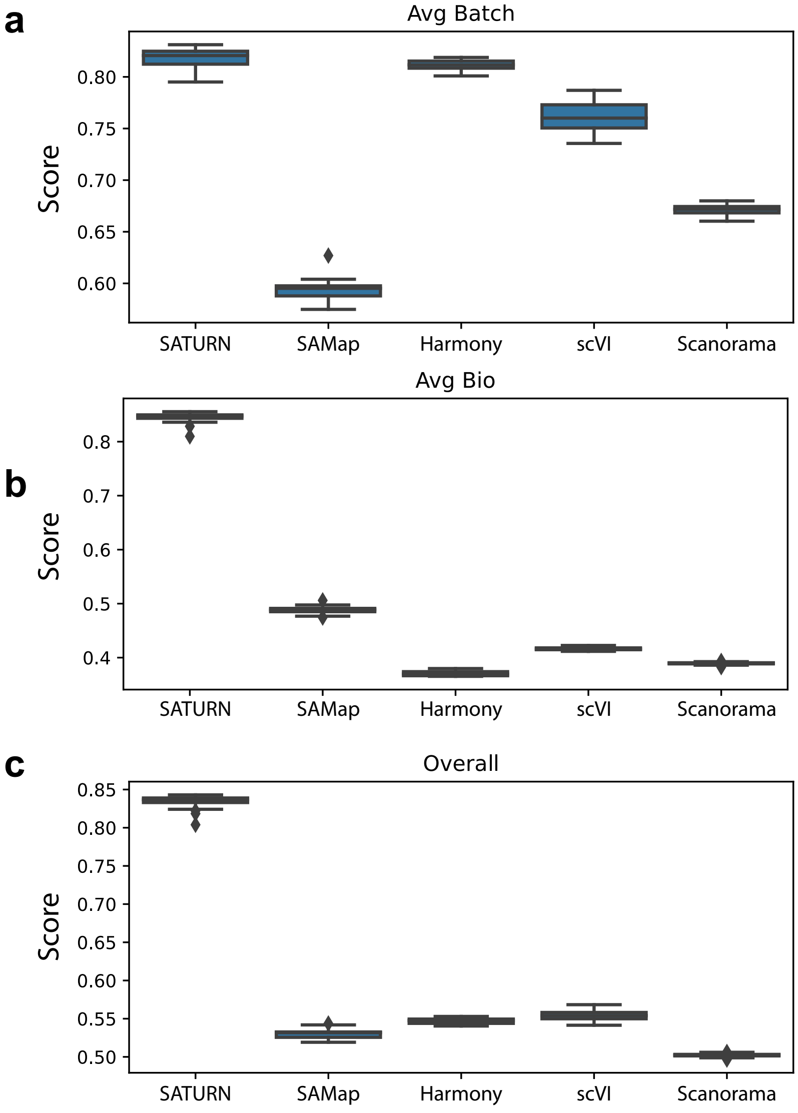


**Supplementary Figure 8: Performance comparison using batch integration evaluation metrics [13].** Median performance of SATURN and baseline methods evaluated using **(a)** weighted mean of the batch removal score (Avg Batch), **(b)** bio-conservation score (Avg Bio), and **(c)** overall score. Distribution is estimated with n = 30 runs of each method. Overall score is calculated as (0.6 * Avg Bio) + (0.4 * Avg Batch). Avg Bio is calculated as the average of NMI cell type score, ARI cell type score, and ASW cell type scores. Avg Batch is calculated as the average of ASW batch score, and graph connectivity scores, where species is taken as the batch variable. Neighbor calculation is done using default Scanpy settings, using each methods' embeddings. Score calculation is done using default SCIB settings [13].

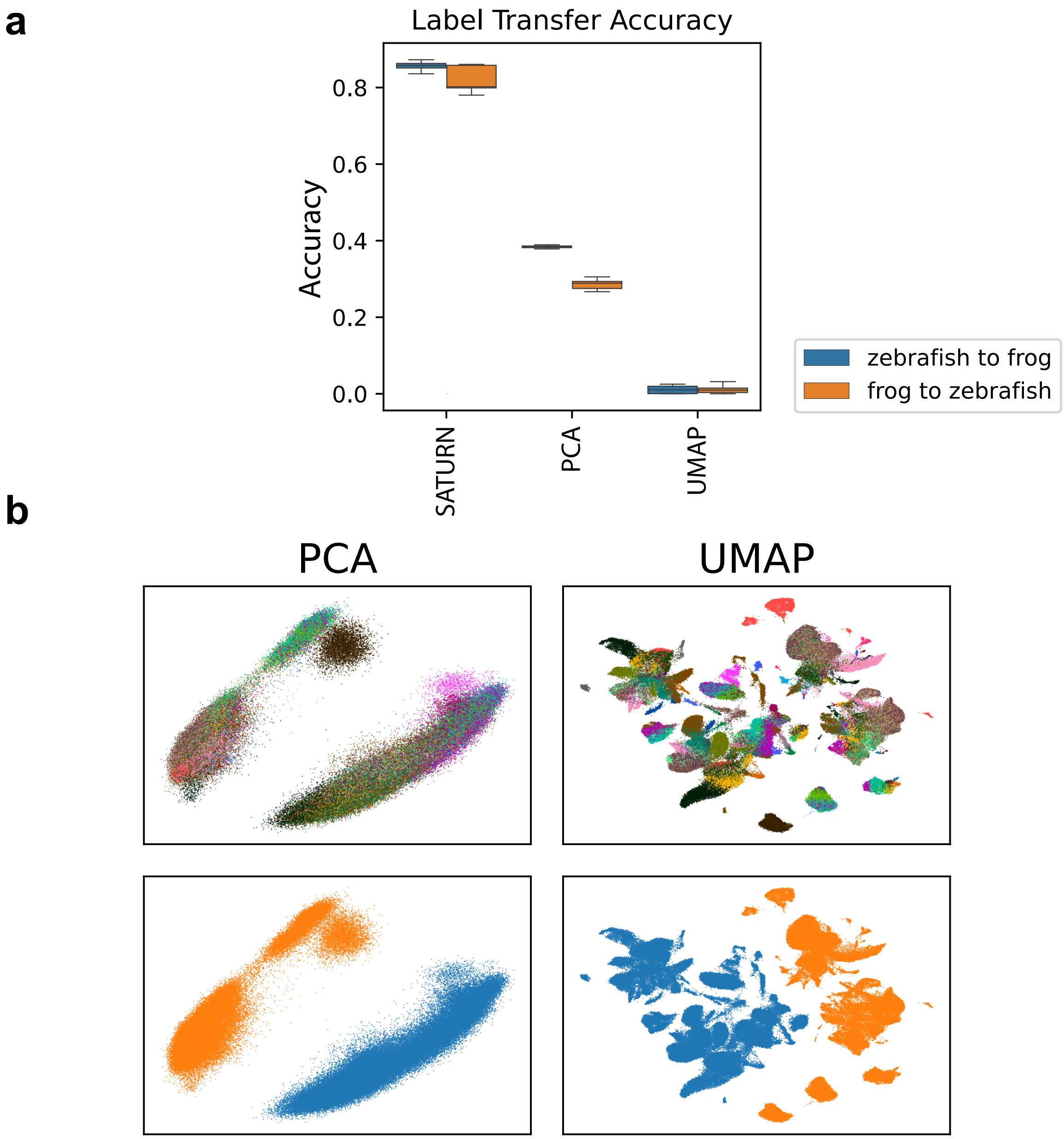


**Supplementary Figure 9: SATURN outperforms UMAP and PCA for cross species integration. (a)** Performance comparison of SATURN versus PCA and UMAP on frog and zebrafish embryogenesis datasets. PCA is calculated using the one-to-one homolog genes as determined by BLAST, followed by expression log normalization. UMAP is then calculated using those top 50 principal components. **B** Visualization of PCA (left) and UMAP (right) embeddings by cell type (top) and species (bottom). For PCA, the top two principal components are used.

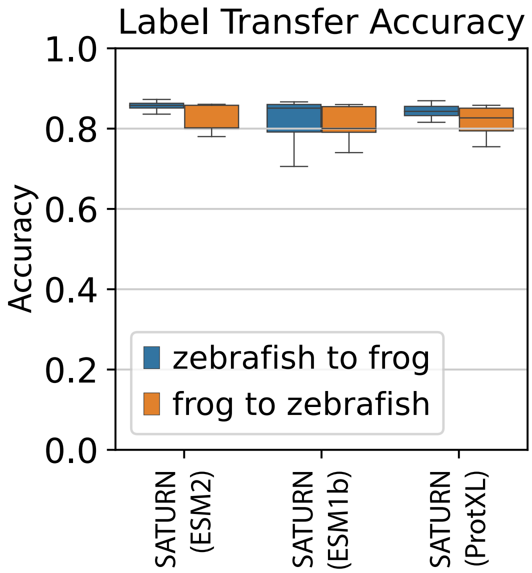


**Supplementary Figure 10: SATURN is robust to protein language model choice.** Median performance of SATURN with different protein language model embeddings on label transfer between frog and zebrafish embryogenesis datasets evaluated using accuracy. Blue boxplots show zebrafish to frog label transfer performance, while orange boxplots show frog to zebrafish label transfer performance. Distribution is estimated with $n = 30$ runs. ESM2 refers to the *esm2 t48 15B UR50D* model [14]. ESM1b refers to the *esm1b t33 650M UR50S* model [15]. ProtXL refers to the *ProtT5 XL U50* model [16].

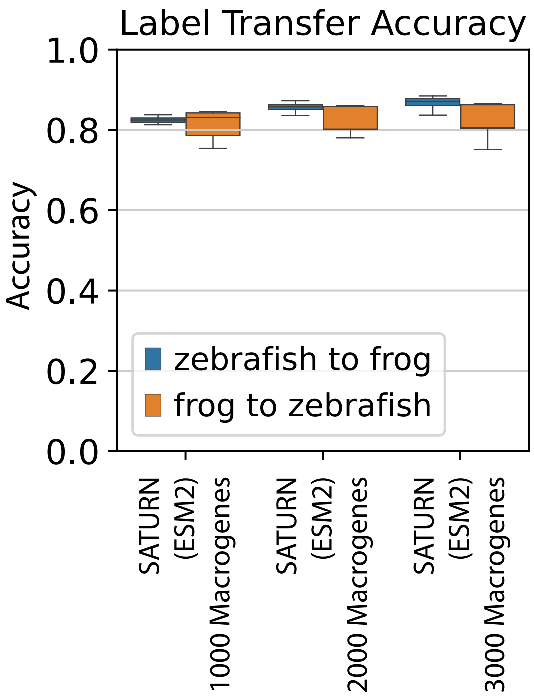


**Supplementary Figure 11: SATURN is robust to choice of number of macrogenes.** Median performance of SATURN with different number of macrogenes on label transfer between frog and zebrafish embryogenesis datasets evaluated using accuracy. Blue boxplots show zebrafish to frog label transfer performance, while orange boxplots show frog to zebrafish label transfer performance. Distribution is estimated with $n = 30$ runs.

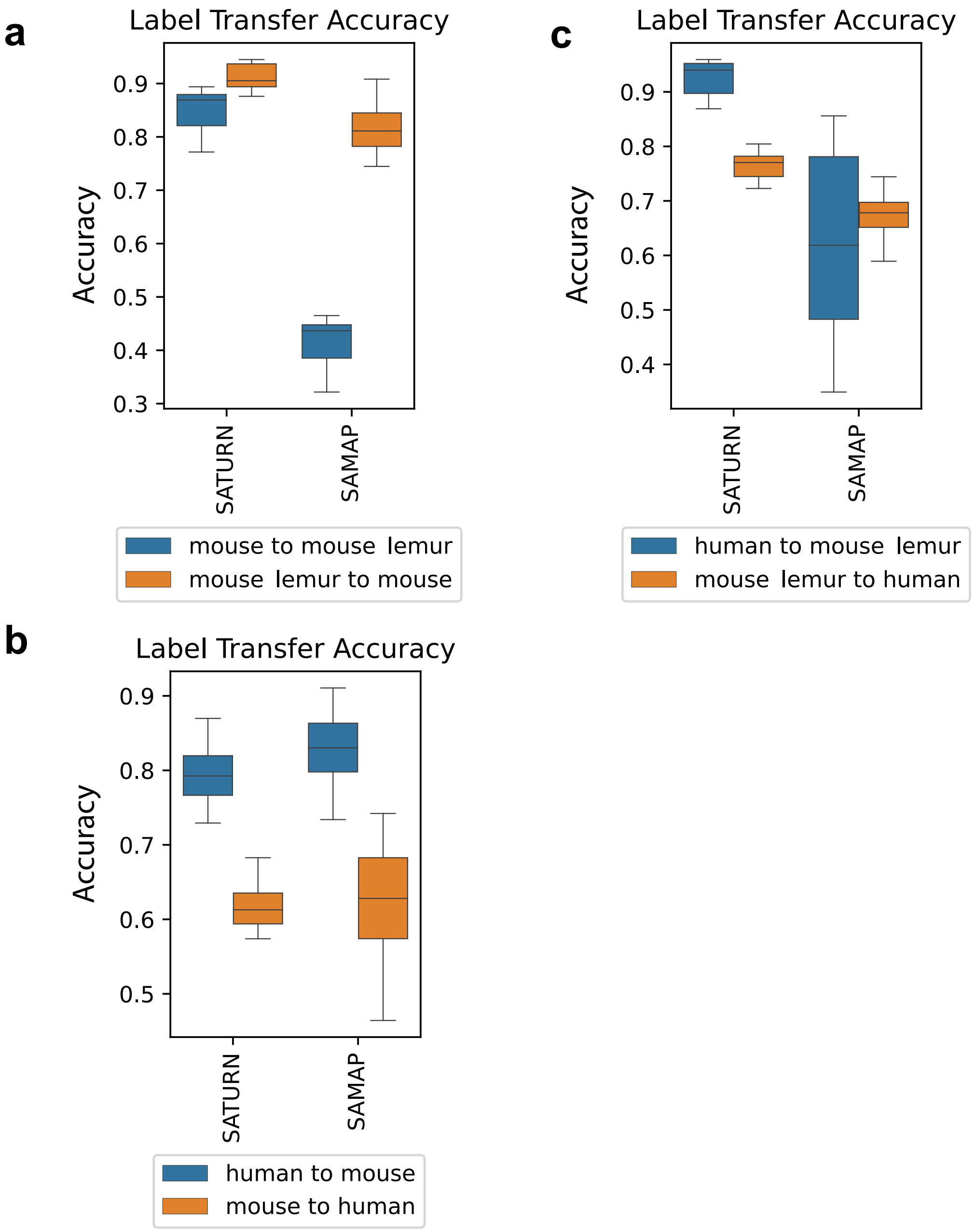


**Supplementary Figure 12: Performance of SATURN and the second best baseline SAMap on transferring annotations on the mammalian cell atlas.** Performance is evaluated using the prediction accuracy of a logistic classifier model trained to differentiate cell types of one species and tested on predicting the cell type annotations of another species. Higher values indicate better performance. SAMap represents a version of the SAMap method in which cell-type annotations are used to integrate datasets. The distribution is obtained with n=30 runs for each method. Performance when transferring annotations from **(a)** mouse to mouse lemur, **(b)** human to mouse, and **(c)** human to mouse lemur.

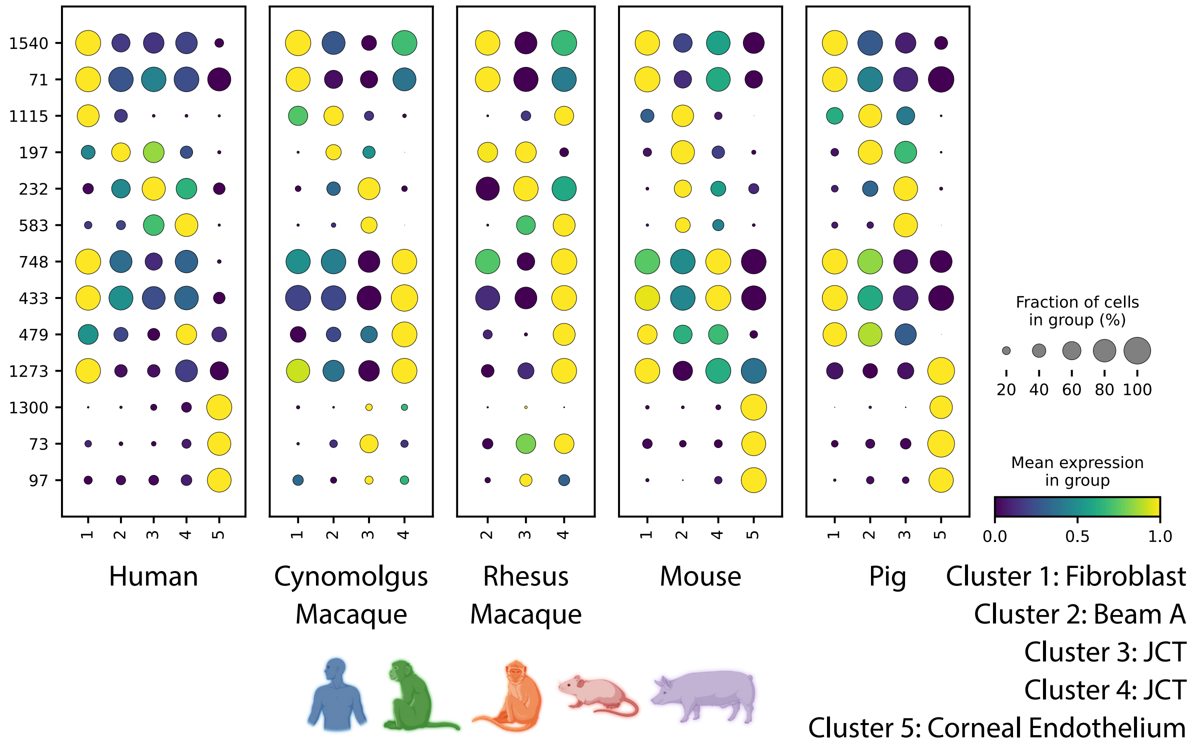


**Supplementary Figure 13: Differentially expressed macrogenes in regrouped AH Atlas cell types.** Rows correspond to macrogene numbers, and columns correspond to cluster numbers. Genes composing each macrogene are listed in Supplementary Table 3. Cluster 1 expresses collagen genes like *Col6a2* (macrogene 1540) which are known fibroblast markers [17]. Cluster 2 expresses *Nr2f1* (macrogene 197) which was identified as a trabecular meshwork marker[17]. Cluster 3 expresses *Rspo* genes (macrogene 583). *Rspo4* was identified as a marker in human JCT [17]. Cluster 4 expresses *Angptl7* (macrogene 479) which was identified as a JCT marker [17]. Cluster 5 expresses corneal endothelium markers including *Ca3* (macrogene 1300). Additionally, Cluster 5 contains a macrogene composed of *Slc4a* genes. Another solute carrier family (SLC) gene [18,19], Slc11a2 was identified as differentially expressed in human corneal endothelium[17].

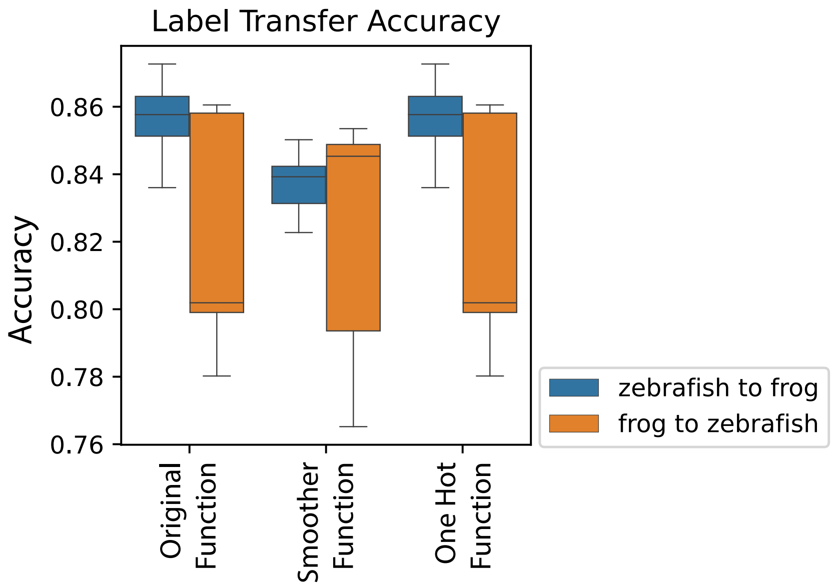


**Supplementary Figure 14: SATURN is robust to choice of macrogene initialization function.** Median performance of SATURN with different macrogene initialization functions evaluated as accuracy of the label transfer between frog and zebrafish embryogenesis datasets. Blue boxplots show zebrafish to frog label transfer performance, while orange boxplots show frog to zebrafish label transfer performance. Distribution is estimated with n = 30 runs.

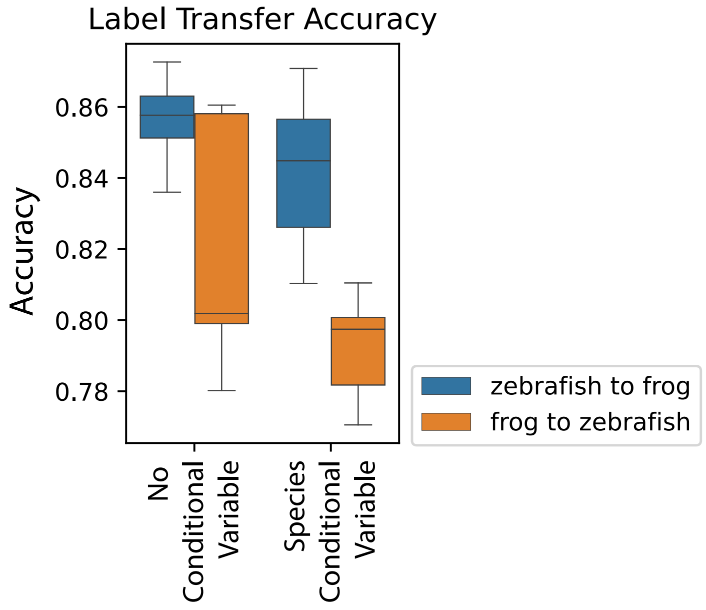


**Supplementary Figure 15: Conditional species variable does not improve performance.** Performance of SATURN using a conditional autoencoder during pretraining with a species conditional variable vs a constant variable. The constant variable is appended to the embedding $\mathbf{z}_c$, while in the conditional variable setting, a one hot representation of the species $s$ is concatenated to the embedding. Blue boxplots show zebrafish to frog label transfer performance, while orange boxplots show frog to zebrafish label transfer performance. Distribution is estimated with $n = 30$ runs.

| | | | | | | | | | |
|---|---|---|---|---|---|---|---|---|---|
| **Macrophage and Myleoid Progenitors** | | | | | | | | | |
| **Arhgdi** | | **Cebp** | | **Ptp** | | **Cybb** | | **Lcp1** | |
| **gene** | **weight** | **gene** | **weight** | **gene** | **weight** | **gene** | **weight** | **gene** | **weight** |
| frog_arhgdib | 2.0586648 | frog_cebpd | 1.7377852 | frog_ptprc | 1.0184758 | frog_cybb | 1.4121561 | frog_lcp1 | 1.1437993 |
| zebrafish_arhgdig | 1.1892534 | frog_cebpb | 1.2713358 | zebrafish_ptprc | 1.0062823 | zebrafish_cybb | 0.98110956 | zebrafish_parvg | 0.95570034 |
| frog_arhgdig | 0.82862365 | zebrafish_cebpa | 1.2589424 | frog_iqcd | 1.0023459 | frog_nox4 | 0.69433695 | zebrafish_parvb | 0.8877349 |
| frog_arhgdia | 0.42018056 | zebrafish_cebpb | 1.1590556 | frog_ptpn6 | 0.975968 | frog_nox1 | 0.6060228 | frog_parva | 0.8088149 |
| frog_c20orf27 | 0.012586672 | frog_mafb | 0.777314 | zebrafish_ptpreb | 0.95713043 | frog_nox5 | 0.016699424 | zebrafish_lcp1 | 0.7965079 |
| frog_arr3 | 0.011929505 | zebrafish_cebpd | 0.6585815 | zebrafish_ptpn6 | 0.9493796 | frog_nadk | 0.012460865 | frog_parvb | 0.7859018 |
| zebrafish_abracl | 0.0018043627 | frog_cebpa | 0.49666032 | zebrafish_ptpn22 | 0.88780427 | zebrafish_slc7a8a | 0.008461936 | frog_parvg | 0.6744168 |
| zebrafish_c7h20orf27 | 0.0017385085 | zebrafish_mafbb | 0.3763802 | zebrafish_ptpn11b | 0.5531652 | frog_rac2 | 0.006649947 | zebrafish_parvab | 0.62013125 |
| zebrafish_arr3b | 0.0011982815 | zebrafish_cebp1 | 0.2561873 | zebrafish_ptprr | 0.533596 | zebrafish_slc7a7 | 0.006204006 | zebrafish_gas2l2 | 0.2786125 |
| zebrafish_cst14b.1 | 0.0007732745 | frog_maf | 0.2419157 | frog_ptprh | 0.39319068 | frog_tfb1m | 0.006183132 | zebrafish_tagln | 0.11108811 |
| **Ionocytes** | | | | | | | | | |
| **Foxi** | | **Dmrt2** | | **Cldn** | | **Ubp1** | | **Atp60v** | |
| **gene** | **weight** | **gene** | **weight** | **gene** | **weight** | **gene** | **weight** | **gene** | **weight** |
| frog_foxi1 | 1.7946975 | frog_dmrt2 | 1.1082501 | zebrafish_cldna | 0.47487783 | frog_ubp1 | 1.4286531 | frog_atp6v0c | 1.5118276 |
| zebrafish_foxi3a | 1.7876347 | zebrafish_dmrt2a | 1.0135943 | zebrafish_cldnh | 0.46664542 | frog_grhl3 | 1.1210046 | zebrafish_atp6v0cb | 1.0511838 |
| zebrafish_foxi1 | 1.7752392 | frog_kank1 | 0.9281069 | frog_cldn4 | 0.45497242 | frog_grhl1 | 1.0977421 | zebrafish_atp6v0ca | 0.6842436 |
| zebrafish_foxg1a | 1.767095 | zebrafish_gcm2 | 0.4140955 | zebrafish_cldnb | 0.40522912 | zebrafish_grhl3 | 1.03748 | frog_atp6v0b | 0.33991408 |
| frog_foxg1 | 1.6482608 | zebrafish_dmrt2b | 0.3073387 | zebrafish_cldne | 0.38158783 | zebrafish_grhl2a | 0.85601187 | zebrafish_atp6v0b | 0.18918027 |
| frog_foxi4.2 | 1.6366866 | zebrafish_cxxc4 | 0.14239863 | zebrafish_lhfpl3 | 0.30014035 | zebrafish_tp63 | 0.6001469 | frog_cnih1 | 0.0017733343 |
| frog_foxi2 | 1.598392 | frog_cxxc4 | 0.1175361 | zebrafish_cldnc | 0.24535778 | frog_grhl2 | 0.54704624 | frog_sec61g | 0.0010816682 |
| zebrafish_foxg1b | 0.92830807 | frog_foxi1 | 0.09924057 | frog_lhfpl4 | 0.24350967 | zebrafish_grhl1 | 0.44004646 | frog_eif1ax | 0.00058003364 |
| zebrafish_foxh1 | 0.7387778 | zebrafish_skor1b | 0.09714537 | zebrafish_lhfpl5a | 0.2244674 | zebrafish_tfcp2l1 | 0.4264244 | zebrafish_sec61g | 0.00044005408 |
| frog_foxe1 | 0.48347607 | frog_hivep1 | 0.07280374 | zebrafish_lhfpl5b | 0.18170683 | frog_tp63 | 0.30589458 | zebrafish_rpl34 | 0.0004284728 |

**Supplementary Table 1: Frog and Zebrafish differentially expressed macrogenes' gene to macrogene weights.** Gene to macrogene weights for the top 10 genes for each differentially expressed macrogene in Figure 2b. Genes are listed in descending order by weight.

| Frog Cell Type | Zebrafish Cell Type | # of Frog Cells | # of Zebrafish Cells | Total # of Cells |
|---|---|---|---|---|
| Hindbrain | Hindbrain | 7273 | 9399 | 16672 |
| Intermediate mesoderm | Intermediate mesoderm | 10324 | 3120 | 13444 |
| Forebrain/midbrain | Forebrain/midbrain | 2081 | 10500 | 12581 |
| Epidermal progenitor | Epidermal progenitor | 9149 | 1921 | 11070 |
| Non-neural ectoderm | Non-neural ectoderm | 8022 | 2227 | 10249 |
| Neural crest | Neural crest | 8393 | 1769 | 10162 |
| Neuroectoderm | Neuroectoderm | 6590 | 3381 | 9971 |
| Placodal area | Placodal area | 6918 | 1188 | 8106 |
| Presomitic mesoderm | Presomitic mesoderm | 6293 | 1642 | 7935 |
| Skeletal muscle | Skeletal muscle | 5772 | 651 | 6423 |
| Neuron | Neuron | 1899 | 4032 | 5931 |
| Tailbud | Tailbud | 1860 | 3759 | 5619 |
| Optic | Optic | 1475 | 3676 | 5151 |
| Blood | Blood | 1569 | 3067 | 4636 |
|  | Pluripotent | 0 | 4277 | 4277 |
| Involuting marginal zone | Involuting marginal zone | 2385 | 1849 | 4234 |
| Endoderm | Endoderm | 2207 | 890 | 3097 |
| Eye primordium | Eye primordium | 2477 | 223 | 2700 |
| Endothelial | Endothelial | 1002 | 884 | 1886 |
| Goblet cell |  | 1473 | 0 | 1473 |
| Small secretory cells |  | 1335 | 0 | 1335 |
| Ionocyte | Ionocyte | 1030 | 292 | 1322 |
| Notochord | Notochord | 766 | 351 | 1117 |
| Blastula |  | 1116 | 0 | 1116 |
| Otic placode | Otic placode | 813 | 270 | 1083 |
| Heart | Heart | 121 | 851 | 972 |
| Spemann organizer |  | 963 | 0 | 963 |
| Myeloid progenitors |  | 778 | 0 | 778 |
| Pronephric mesenchyme |  | 777 | 0 | 777 |
| Cement gland primordium |  | 721 | 0 | 721 |
| Lens | Lens | 458 | 210 | 668 |
|  | Rare epidermal subtypes | 0 | 513 | 513 |
| Notoplate | Notoplate | 339 | 115 | 454 |
| Rohon-beard neuron | Rohon-beard neuron | 134 | 289 | 423 |
| Olfactory placode | Olfactory placode | 139 | 276 | 415 |
|  | Macrophage | 0 | 405 | 405 |
|  | Periderm | 0 | 382 | 382 |
| Hatching gland | Hatching gland | 180 | 82 | 262 |
|  | Dorsal organizer | 0 | 233 | 233 |
|  | Pharyngeal pouch | 0 | 209 | 209 |
|  | Apoptotic-like | 0 | 163 | 163 |
|  | Pronephric duct | 0 | 95 | 95 |
| Germline | Germline | 33 | 53 | 86 |
| Neuroendocrine cell |  | 70 | 0 | 70 |
|  | Pancreas primordium | 0 | 49 | 49 |
|  | Secretory epidermal | 0 | 34 | 34 |
|  | Apoptotic-like 2 | 0 | 33 | 33 |
|  | Forerunner cells | 0 | 5 | 5 |
|  | Epiphysis | 0 | 3 | 3 |
|  | Nanog-high | 0 | 3 | 3 |
| Totals: 36 | 42 | 96935 | 63371 | 160306 |

**Supplementary Table 2: Cell Type Matching and Frequencies in Frog and Zebrafish Embryogenesis.** Cell type pairs used for scoring frog and zebrafish embryogenesis embeddings, and cell type counts.

| Cluster | Macrogene | Human Genes | Cynomologus Macaque Genes | Rhesus Macaque Genes | Mouse Genes | Pig Genes |
|---|---|---|---|---|---|---|
| 1 | 1540 | Col6A2, Vit, Col6A6 | Vit, Col28A1, Antxr2 | Col6A2, Col28A1, Vit | Col6A1, Col6A2, Vit | Vit, Antxr2, Col6A2 |
| 1 | 71 | Rpp25, Sco2, Siglec1 | Adam15, Siglec1, Nop9 | Adam15, Lhb, Kcp | Ptpn18 | C4A, Kcnk7, Rpp25 |
| 2 | 1115 | Cxcl12, Ccl25 | Cxcl12 | Ccl25, Cxcl14 | Cxcl12 | Cxcl12 |
| 2 | 197 | Nr2F1, Nr2F2 | Nr2F1, Nr2E3, Nr2E1 | Nr2F2, Nr2F1, Nr2E3 | Nr2F1, Nr2F2, Nr2E3 | Nr2F1, Nr2F2, Nr0B2 |
| 3 | 232 | Tagln, Tagln2, Tagln3 | Tagln, Tagln3 | Tagln, Tagln2 | Tagln, Tagln3 | Tagln, Tagln3 |
| 3 | 583 | Rspo2, Rspo3 | Rspo2, Rspo3 | Rspo2, Rspo3 | Rspo3, Rspo2, Rspo1 | Rspo3, Rspo2 |
| 4 | 748 | Bgn | | | | |
| 4 | 433 | Prelp, Ogn, Aspn | Ogn, Kera, Prelp | Ogn, Prelp, Optc | Dcn, Fmod, Optc | Omd, Ogn, Ecm2 |
| 4 | 479 | Angptl7, Fgl2, Angptl1 | Fgl2, Angptl7, Fgb | Fgl2, Angptl7, Fgg | Fgl2, Angptl7, Angptl2 | Fgl2, Angptl7, Fibcd1 |
| 4 | 1273 | Tnxb, Matn2, Tnr | Tnc, Morn4 | Morn4, Tnc, Matn2 | Tnxb | Zcchc13, Tnc, Tnr |
| 5 | 1300 | Ca3, Ca13, Ca7 | Ca2, Ca7, Ca3 | Ca3, Ca2, Ca13 | Car3, Car2, Car13 | Ca2, Ca3, Ca7 |
| 5 | 73 | Slc4A7, Slc4A4, Slc4A10 | Slc4A10, Slc4A7, Slc4A4 | Slc4A10, Slc4A7, Slc4A4 | Slc4A4, Slc4A5 | Slc4A4, Slc4A7, Slc4A5 |
| 5 | 97 | Fgf6, Fgf23, Fgf16 | Fgf21, Fgf19, Fgf10 | Fgf10, Fgf8, Fgf9 | Fgf10, Fgf5, Fgf21 | Fgf10, Fgf22, Fgf21 |

**Supplementary Table 3: Differentially expressed macrogenes in regrouped AH Atlas cell types.** Genes in the table represent the corresponding species' top 3 genes per macrogene, ordered by weight and with weights above $0.5$.
